# Integrated analysis reveals FOXA1 and Ku70/Ku80 as direct targets of ivermectin in prostate cancer

**DOI:** 10.1101/2022.01.19.476914

**Authors:** Shidong Lv, Zeyu Wu, Mayao Luo, Yifan Zhang, Jianqiang Zhang, Laura E. Pascal, Zhou Wang, Qiang Wei

## Abstract

Ivermectin is a widely used antiparasitic drug and shows promising anticancer activity in various cancer types. Although multiple signaling pathways modulated by ivermectin have been identified, few studies have focused on the exact target of ivermectin. Herein, we report the pharmacological effects and direct targets of ivermectin in prostate cancer (PCa). Ivermectin caused G0/G1 arrest, induced cell apoptosis, DNA damage, and decreased androgen receptor (AR) signaling in PCa cells. Using integrated omics profiling, including RNA-seq and thermal proteome profiling, we found that the forkhead box protein A1 (FOXA1) and non-homologous end joining (NHEJ) repair executer Ku70/Ku80 were the direct targets of ivermectin. The binding of ivermectin and FOXA1 reduced the chromatin accessibility of AR and the G0/G1 cell cycle regulator E2F1, leading to cell proliferation inhibition. The binding of ivermectin and Ku70/Ku80 impaired the NHEJ repair ability. Cooperating with the downregulation of homologous recombination repair after AR inhibition, ivermectin triggered synthetic lethality. Our findings demonstrate the anticancer effect of ivermectin in prostate cancer, indicating that its use may be a new therapeutic approach for PCa.

## Introduction

Prostate cancer is the most frequently diagnosed cancer among men and ranks as the second leading cause of cancer-related deaths in the United States of America, with more than 240,000 diagnoses and over 34,000 deaths annually(Siegel *et al*, 2021). With surgical resection, in combination with androgen deprivation treatment (ADT) when necessary, the 5-year survival rate of early-stage prostate cancer is 98%. However, once the disease has progressed to castration-resistant prostate cancer (CRPC), the survival duration is only 1–2 years on average(Halabi *et al*, 2016). Due to androgen receptor (AR) overexpression, mutation, and splice variants, AR can be re-activated, resulting in resistance to current anti-androgen drugs(Carceles-Cordon *et al*, 2020). Genetic alterations of AR have been reported in up to 57.78% of advanced prostate cancer cases(Abida *et al*, 2019). Despite several strategies that have been proposed to improve this situation, the prognosis for patients with CRPC remains poor(Davis *et al*, 2019; Rathkopf *et al*, 2014), thereby highlighting the need to develop new therapeutic agents/approaches.

Drug repositioning is a highly studied alternative strategy for the discovery and development of anticancer drugs. This strategy identifies new indications for existing pharmacological compounds. Ivermectin is a macrolide antiparasitic drug with a 16-membered ring derived from avermectin(Campbell *et al*, 1983), which was approved by the Food and Drug Administration (FDA) for the treatment of onchocerciasis in humans in 1978(Laing *et al*, 2017). To date, ivermectin has been used by millions of people worldwide and exhibits a wide margin of clinical safety(Juarez *et al*, 2018a). Recently, several studies have explored the potential of ivermectin as a new cancer treatment(Crump, 2017; Juarez *et al*, 2018b; Tang *et al*, 2020). In breast cancer, ivermectin decreases p21-activated kinase 1 (PAK1) expression by promoting its degradation and inducing cell autophagy(Dou *et al*, 2016). In ovarian cancer, ivermectin can block the cell cycle and induce cell apoptosis through a Karyopherin-β1 (KPNB1) related mechanism(Kodama *et al*, 2017). In leukemia, ivermectin preferentially kills leukemia cells at low concentrations by increasing the influx of chloride ions into cells, which trigger plasma membrane hyperpolarization and reactive oxygen species (ROS) production(Sharmeen *et al*, 2010). These results not only confirm the promising effect of ivermectin, but also reveal its safety for tumor suppression through the *in vivo* analysis. However, the detailed mechanism and direct target of ivermectin underlying ivermectin-mediated tumor suppression remain to be further elucidated.

Here, we showed that ivermectin suppresses prostate cancer progression efficiently both *in vitro* and *in vivo*. We applied integrated profiling including RNA-seq and Thermal proteome, that found pioneer factor Forkhead Box Protein A1 (FOXA1) and Non-homologous End Joining (NHEJ) repair executer Ku70/Ku80 was the direct target of Ivermectin in prostate cancer. Ivermectin binds to these two proteins and blocks their biological function, which results in blockade of AR signaling transcription, E2F1 expression, and deficiency of DNA double-strand break (DSB) repair system, and thereby leads to G0/G1 arrest and trigger synthetic lethality. Our findings demonstrate both the effect and target of ivermectin in prostate cancer comprehensively and systemically, indicating that the use of ivermectin may constitute a new therapeutic approach for prostate cancer.

## Results

### Ivermectin preferentially inhibited the viability of AR-positive prostate cancer cells

To evaluate the effect of ivermectin in prostate cancer, we analyzed cell viability using MTT assays in AR-positive prostate cancer cell lines, LNCaP, C4-2, and 22RV1, AR-negative prostate cancer cell lines DU145 and PC-3, and non-tumorigenic human prostate primary stromal cells from patients with BPH(Chen *et al*, 2020). As is shown in **Fig. 1**, ivermectin markedly decreased the viability of all prostate cancer cells in a dose-dependent manner. Compared to tumor cells, the IC50 of ivermectin in primary BPH stromal cells was much higher. Moreover, the effect of ivermectin was more dramatic in AR-positive prostate cancer cells than in AR-negative prostate cancer cells. The IC50 value was 2–3-fold lower in LNCaP and C4-2 cells than in DU145 and PC-3 cells. Meantime, the 22RV1 also showed dramatic responsive to ivermectin, suggesting that AR variants did not compromise the effect of ivermectin. Overall, our data revealed that ivermectin exerted a profound suppression of prostate cancer across different stages of the disease.

**Figure 1.**
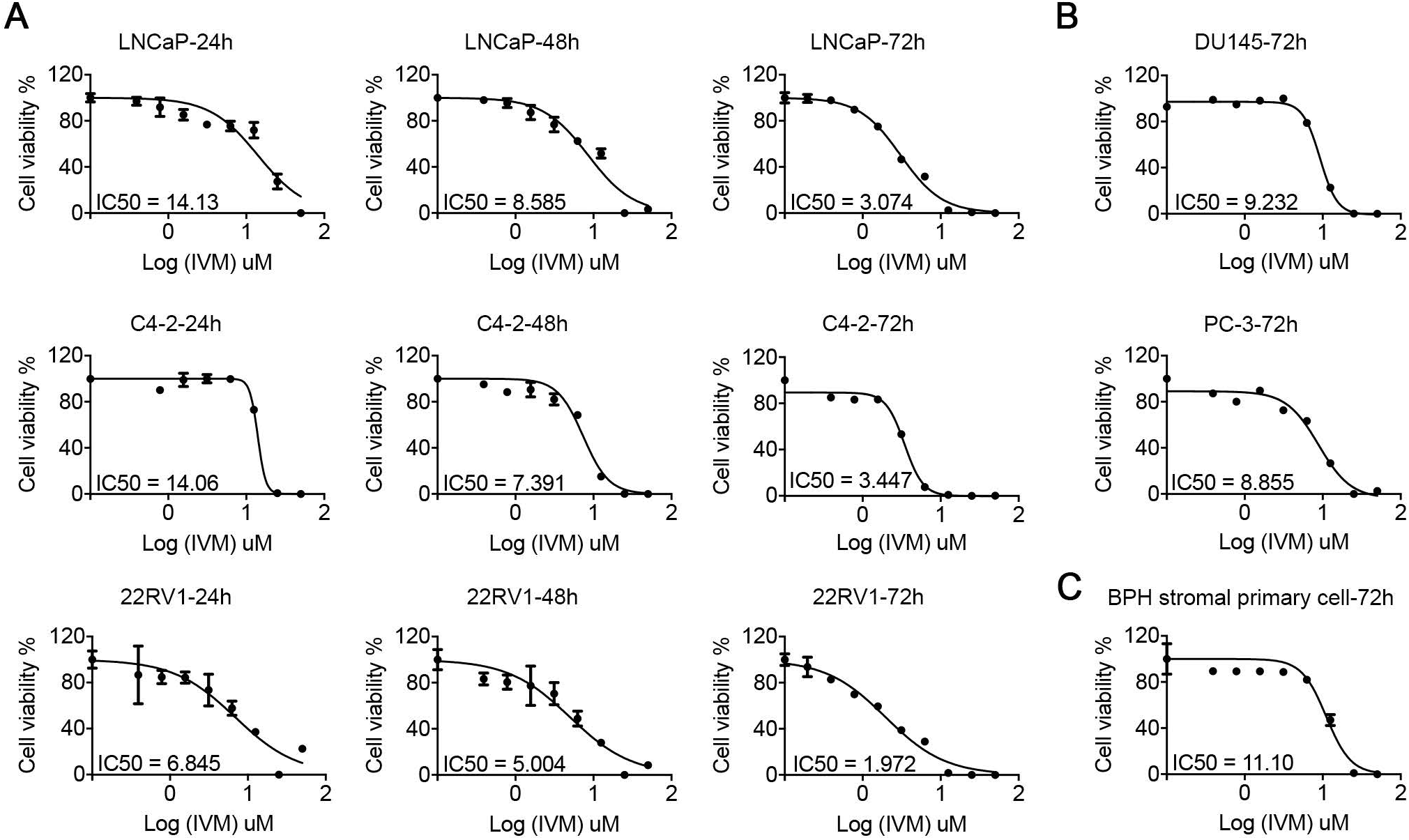
Ivermectin inhibited prostate cancer cell viability. Cell viability was measured by the MTT assay in AR positive cells (LNCaP, C4-2, and 22RV1, **A**), AR negative cells (DU145 and PC-3, **B**), and prostate primary cells from benign prostatic hyperplasia patients (**C**) treated with the indicated concentrations of ivermectin for either 24 h, 48 h, or 72 h.

### Ivermectin induced G0/G1 arrest, apoptosis, and DNA damage in prostate cancer cells

To further address ivermectin inhibition in prostate cancer cells, we explored the cell cycle distribution in response to ivermectin using flow cytometry. Consistent with the cell viability results, an ivermectin treatment of 48 h significantly arrested the cell cycle at the G0/G1 phase in LNCaP, C4-2, and 22RV1 cells (**Fig. 2A)**. Meanwhile, in the high-dose group (12 μM), we observed marked sub-G1 peaks in C4-2 and 22RV1 cells, indicating that ivermectin could induce cell apoptosis (**Supplementary Fig. S1A**). Thus, we further explored the cell apoptosis rate after the ivermectin treatment using PI/annexin V staining. As expected, a high-dose ivermectin treatment for 48 h significantly induced apoptosis in LNCaP, C4-2, and 22RV1 cells (**Fig. 2B**). In line with this, an obvious upregulation of apoptosis markers, cleaved PARP and cleaved caspase-3, was detected in ivermectin-treated cells (**Fig. 2C**).

**Figure 2.**
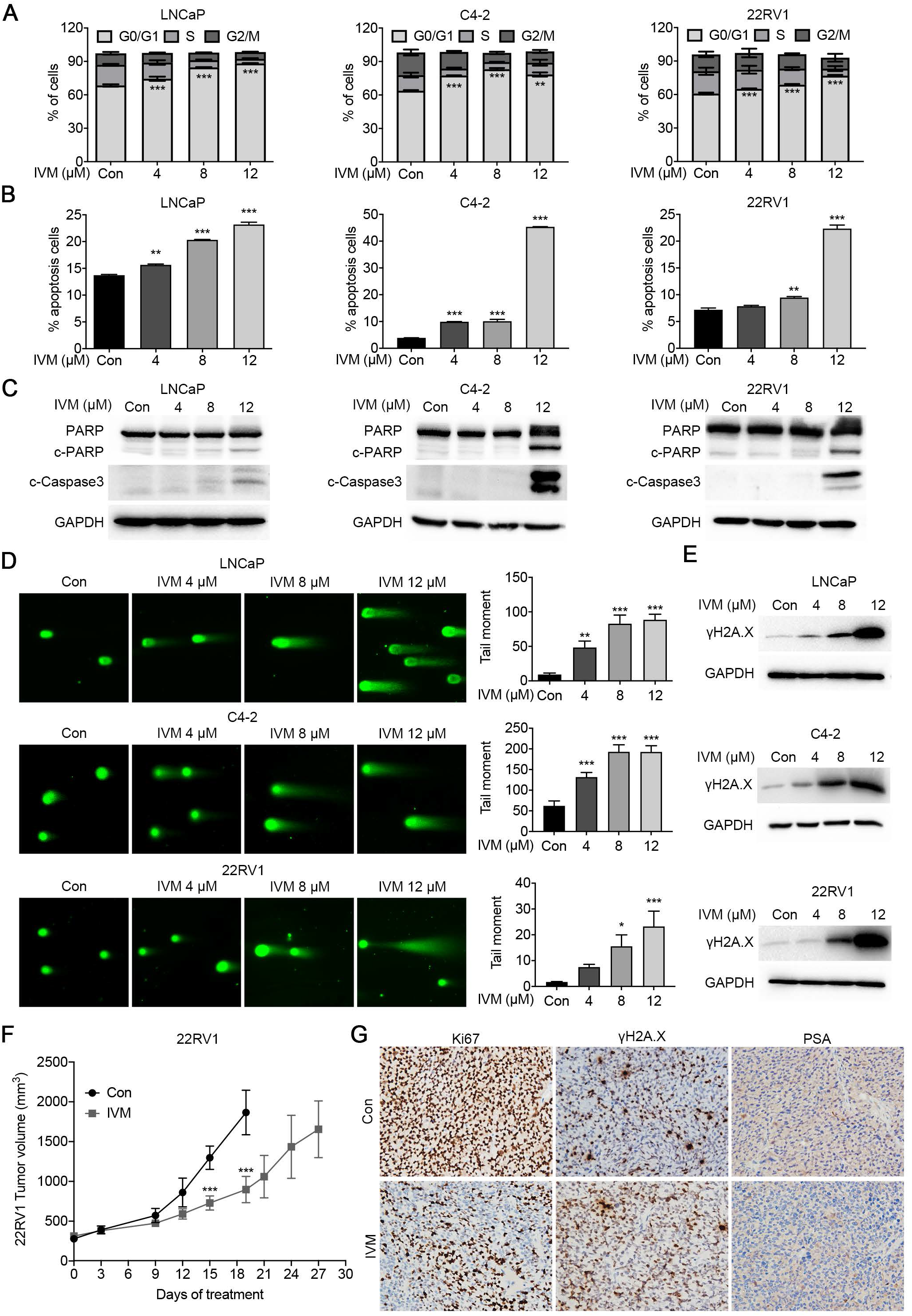
Ivermectin led to G0/G1 arrest, apoptosis, and DNA damage in prostate cancer. **(A)** The ivermectin arrest cell cycle at G0/G1 was measured by flow cytometry. LNCaP, C4-2, and 22RV1 cells were treated with ivermectin at 4 μM, 8 μM, and 12 μM for 48 h. **(B)** Ivermectin induced cell apoptosis detected by PI/Annexin V staining. Cells were treated as in A. The PI+/Annexin V+ and PI-/Annexin V+ cells were calculated as apoptotic cells. **(C)** Western blot analysis of PARP and cleaved-Caspase3 (c-Caspase3) in cells treated with ivermectin for 48 h. **(D)** Ivermectin increased DNA damage. DNA fragments were shown as comet images in alkaline gel electrophoresis. The tail moment was used to quantify the DNA damage in the treatment of ivermectin for 48 h. **(E)** Western blot analysis of γH2A.X in cells treated with the ivermectin for 48 h. **(F)** Tumor volume of 22RV1 xenografts after castration treated with vehicle (con) or ivermectin (10 mg/kg, n = 5 for each group). **(G)** Representative images of Ki67, γH2A.X and PSA immunostaining, in 22RV1 tumors treated with vehicle or ivermectin.

Increased DNA damage is one of the most common characteristics of anticancer drugs. We used comet assay to evaluate DNA damage levels after the ivermectin treatment. As is shown in **Fig. 2D**, the comet assay moment increased dramatically in a dose-dependent manner in ivermectin-treated LNCaP, C4-2, and 22RV1 cells. Moreover, an elevated expression of the DNA damage marker γH2A.X was observed after the ivermectin treatment in all three cell lines (**Fig. 2D**). DNA damage activates DNA damage response proteins, leading to senescence and apoptosis(Lv *et al*, 2019). To understand better the cell fate after the ivermectin treatment, we also assayed cell senescence by β-galactosidase staining. An ivermectin treatment for 48 h had no obvious effect on senescence in any of the tested prostate cancer cell lines, LNCaP, C4-2, and 22RV1 (**Supplementary Fig. S1B**).

Based on the MTT assay, AR-negative PC-3 and DU145 cells were less sensitive to the ivermectin treatment. This observation was confirmed. An ivermectin treatment of 48 h had no significant effect on the cell cycle (**Supplementary Fig. S2A**), or apoptosis in DU145 cells (**Supplementary Fig. S2B**). The comet assay showed that a high-dose ivermectin treatment (12 μM) induced DNA damage, while low and median doses showed no effect (**Supplementary Fig. S2C**).

22RV1 xenograft model was used to determine effect of ivermectin on CRPC progression in *vivo*. Male mice bearing 22RV1 xenografts were castrated when tumors exceeded 300 mm^3^ and randomized to vehicle or Ivermectin administered 10 mg/kg 3 times per week. Ivermectin significantly reduced 22RV1 tumor volume growth (**Fig. 2F**), lowering Ki-67 and PSA levels, and increasing the γH2A.X level in tumor tissue (**Fig. 2G**).

Taken together, these results revealed that ivermectin could inhibit prostate cancer progression *in vitro* and *in vivo* by inducing G0/G1 arrest, apoptosis, and DNA damage.

### Ivermectin inhibited AR signaling in prostate cancer cells

Cell viability and functional assays highlighted the close relationship between ivermectin and the AR signaling pathway. Western blotting showed that ivermectin markedly reduced AR and prostate-specific antigen (PSA) protein expression in LNCaP and C4-2 cells (**Fig. 3A**). Real-time quantitative reverse transcription PCR (RT-qPCR) analysis of AR downstream targets supported the inhibition of the AR signaling pathway by ivermectin (**Fig. 3B**). Moreover, in addition to full-length AR (AR-FL), ivermectin also reduced the expression of AR variants (ARVs) and AR downstream targets in 22Rv1 cells (**Fig. 3C and 3D**). We tested the effect of ivermectin on ARVs in two other cell lines, LN95 and VCaP. Similar to its effect on 22RV1 cells, ivermectin decreased the expression of AR-FL and ARVs, and increased the expression of cleaved-PARP and γH2A.X (**Fig. 3E**), indicating that ivermectin was a competent inhibitor of both AR-FL and ARVs. To further identify the inhibition role of ivermectin on AR signaling pathway, the R1881 induction assay were subsequently performed. As is shown in **Fig. 3F**, ivermectin could compete the increased AR transcription activity after R1881 treatment. Interestingly, the R1881 treatment only partially reversed ivermectin-mediated cell apoptosis and DNA damage (**Fig. 3F**), suggesting that there was an AR-independent pathway for the effect of ivermectin in prostate cancer. This observation was supported by cell cycle analysis. Ivermectin arrested cells at the G0/G1 phase either with or without the R1881 treatment (**Fig. 3G**).

**Figure 3.**
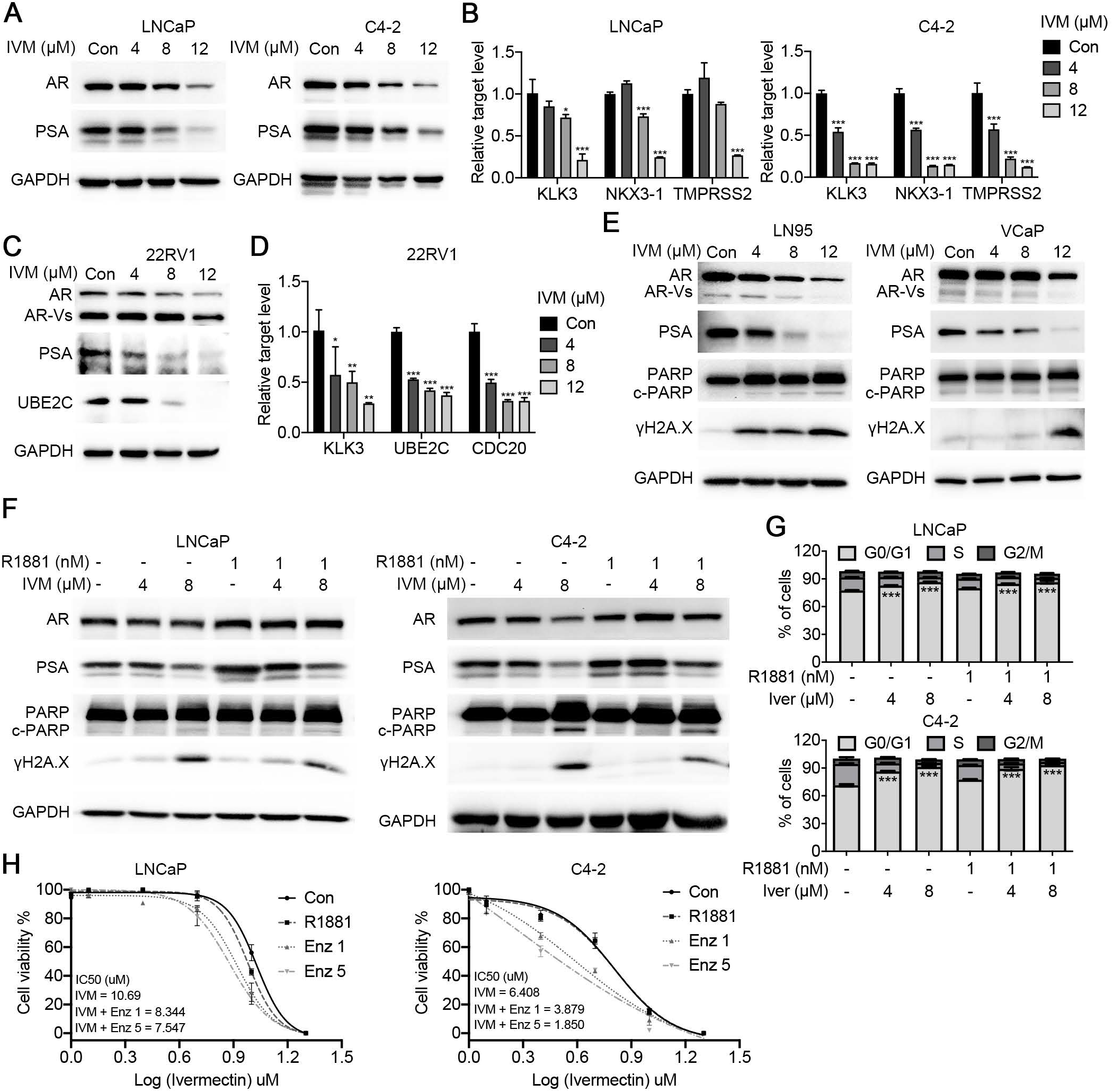
Ivermectin inhibited the FL-AR and AR-V7 signaling activity. **(A)** Western blot analysis of AR and PSA in LNCaP and C4-2 cells treated with ivermectin for 48 h. **(B)** RT-qPCR analysis of AR target genes (*KLK3, TMPRSS2*, and *NKX3-1*) in LNCaP and C4-2 cells treated with ivermectin for 48 h. **(C)** Western blot analysis of FL-AR, ARVs, PSA, and UBE2C in ivermectin-treated 22RV1 cells at 48 h. **(D)** RT-qPCR analysis of KLK3 and ARV target genes (*UBE2C* and *CDC20*) in 22RV1 cells treated with ivermectin for 48 h. **(E)** Western blot analysis of FL-AR, ARVs, PSA, PARP, and γH2A.X in the other two ARV positive cells lines, LN95 and VCaP, treated with ivermectin for 48 h. **(F)** Western blot analysis of AR, PSA, PARP, and γH2A.X in LNCaP and C4-2 cells after the implementation of 4 μM and 8 μM of ivermectin with or without 1 nM R1881. **(G)** Ivermectin inhibited the cell cycle at G0/G1 in the presence of R1881. LNCaP and C4-2 cells were treated with ivermectin at 4 and 8 μM for 48 h in the absence or presence of 1 nM R1881. **(H)** Cell viability was measured by the MTT assay. LNCaP and C4-2 cells were treated with indicated concentrations of ivermectin for 48 h with or without 5 μM and 10 μM enzalutamide for 48 h.

In addition, we tested the combination of ivermectin and enzalutamide. The results showed that the IC50 of ivermectin in the combination treatment group was much lower than that in the ivermectin single drug group (**Fig. 3H**). Thus, the AR-dependent and AR-independent pathways would cooperate with each other for the tumor suppressive role of ivermectin. Together, our data indicate that ivermectin is a novel approach to suppress AR genomic alterations that drive resistance in CRPC.

### Ivermectin downregulated the expression of E2F targets

To further explore the molecular association of ivermectin action in prostate cancer, we characterized the transcriptional profile altered by ivermectin by performing RNA-seq in C4-2 and 22RV1 cells treated with two different doses of ivermectin in regular medium. Consistent with its AR inhibitory effect, ivermectin suppressed the expression of downstream targets of FL-AR (**Supplementary Fig. S3A and S3B**) and ARVs (**Supplementary Fig. S3C**). Further gene set enrichment analysis (GSEA)(Mootha *et al*, 2003) revealed the positive enrichment of hallmark gene sets associated with apoptosis (e.g., apoptosis and the P53 pathway), and the suppression of gene sets related to proliferation, cell cycle, and DNA damage repair (e.g., E2F targets, the mitoticspindle, MYC targets V1/2, the G2M checkpoint, and DNA damage repair; **Fig. 4A**). After combining differentially expressed genes (DEGs) from these two cell lines, a total of 2,997 concordant DEGs were identified (**Fig. 4B**) and the GSEA analysis was repeated. Among all the alterations, the E2F targets constituted the most dramatically and consistently downregulated set in both the C4-2 and 22RV1 cells (**Fig. 4C**). This observation was further confirmed by another database of transcription factor binding sites, TRANSFAC(Kaplun *et al*, 2016) (**Fig. 4D**). Moreover, our results showed that both the protein level (**Fig. 4E**) and mRNA level (**Fig. 4F**) of E2F1 decreased after administering the ivermectin treatment in a dose-dependent manner. E2F1 activity is important to drive the cell cycle from the G1 to the S phase(Fang *et al*, 2020), consisting with our finding in cell functional analysis. To further explore the interaction between ivermectin and E2F1, the CETSA(Jafari *et al*, 2014; Molina *et al*, 2013) was performed. However, we failed to identify the direct binding between ivermectin and E2F1 in C4-2 cells (**Fig. 4G**), indicating E2F1 was not a direct target of ivermectin. Collectively, these data suggested that ivermectin could target other proteins that regulate E2F1 expression at the transcriptional level.

**Figure 4.**
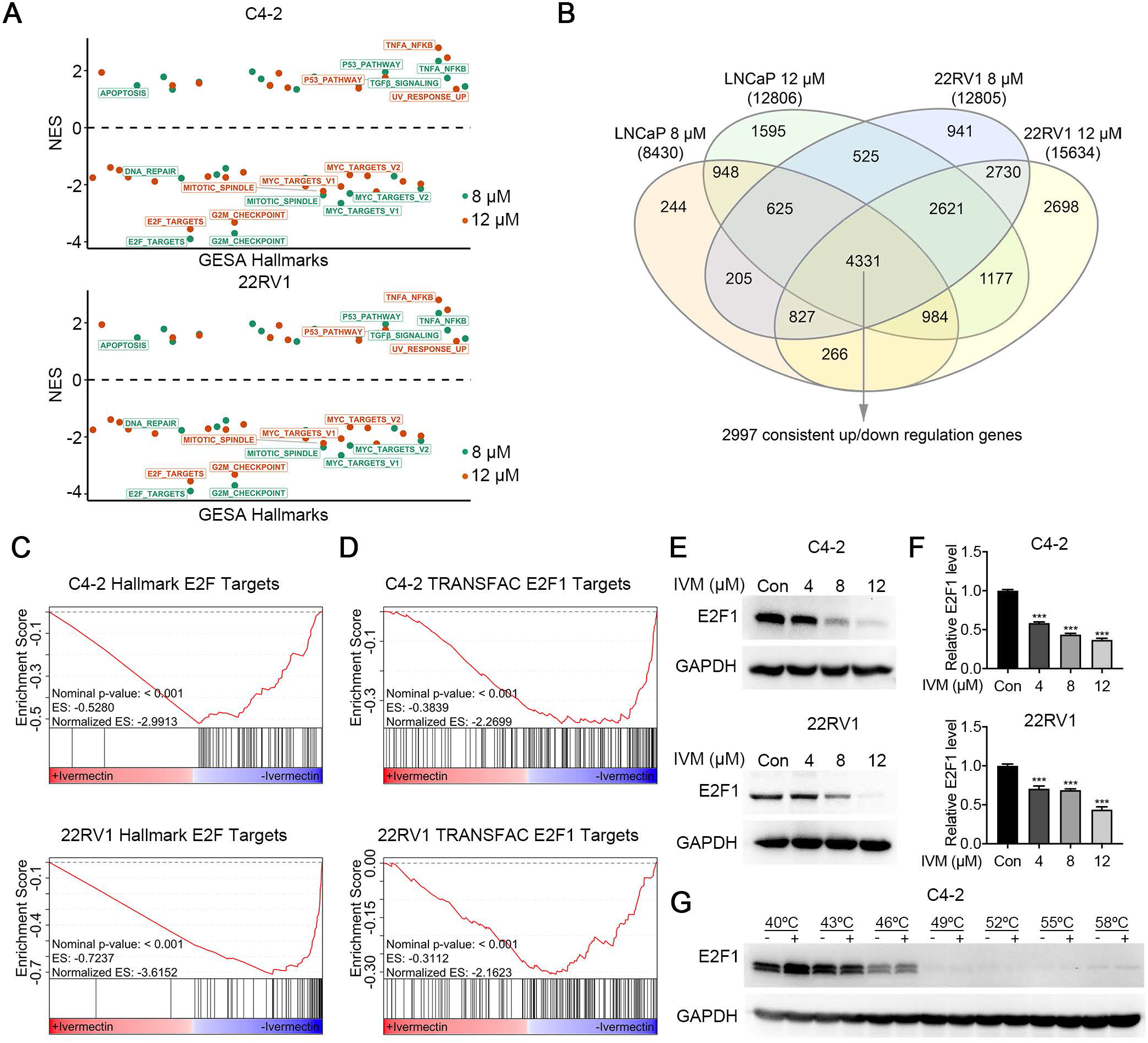
Ivermectin repressed E2F targets. **(A)** Normalized-enrichment scores (NES) of GSEA hallmark gene sets for all four comparation in C4-2 and 22RV1 cells. Significant gene sets comparing ivermectin versus vehicle (P value < 0.05) are labeled. **(B)** Venn diagram indicating the number of DEGs between C4-2 and 22RV1 cells. **(C-D)** The GSEA of C4-2 and 22RV1 concordant altered genes highlighted that hallmark E2F targets **(C)** and TRANSFAC E2F1 targets **(D)** were repressed by ivermectin. **(E-F)** The protein **(E)** and mRNA **(F)** expression of E2F1 decreased in C4-2 and 22RV1 cells treated with ivermectin. **(G)** Western blots showing thermostable E2F1 following indicated heat shocks in the presence (+) or absence (−) of 50 μM ivermectin in C4-2 cells.

### Ivermectin bound and blocked the function of pioneer factor FOXA1

FOXA1 is a pioneer transcription factor that functions to loosen the compact chromatin to facilitate the binding of steroid receptors such as estrogen receptor and AR(Gao *et al*, 2019). A recent study showed that FOXA1 could promote G1 to the S-phase transit by acting as an upstream regulator of E2F1(Zhang *et al*, 2011). These findings, along with the effect of ivermectin on prostate cancer, suggest that FOXA1 is a potential candidate target of ivermectin. To address this, the first step was to analyze the effect of ivermectin on FOXA1 regulated genes. GSEA revealed that genes induced by FOXA1 in the absence of androgens(Jin *et al*, 2013) significantly overlapped with those repressed by ivermectin (**Fig. 5A, left**). This observation was confirmed by RT-qPCR. FOXA1-induced genes decreased significantly in the ivermectin-treated group (**Fig. 5B, left**). However, the alteration of FOXA1-repressed genes was not significant (**Fig. 5A, right**). In contrast to FOXA1-induced genes, FOXA1-repressed genes oppose the action of AR signaling and are reported to correlate with epithelial mesenchymal transformation (EMT)(Jin *et al*., 2013). RT-qPCR showed that the expression of EMT-related genes, including *MET, MMP7*, and *SOX9*, decreased (**Fig. 5B, right**). Moreover, the western blot results showed that the expression of N-cadherin decreased consistently after the ivermectin treatment, while the expression of FOXA1 decreased only slightly (**Fig. 5C**). These results indicate that ivermectin could inhibit FOXA1 signaling activity without promoting cancer metastasis, unlike other drugs targeting FOXA1(Wang *et al*, 2020).

**Figure 5.**
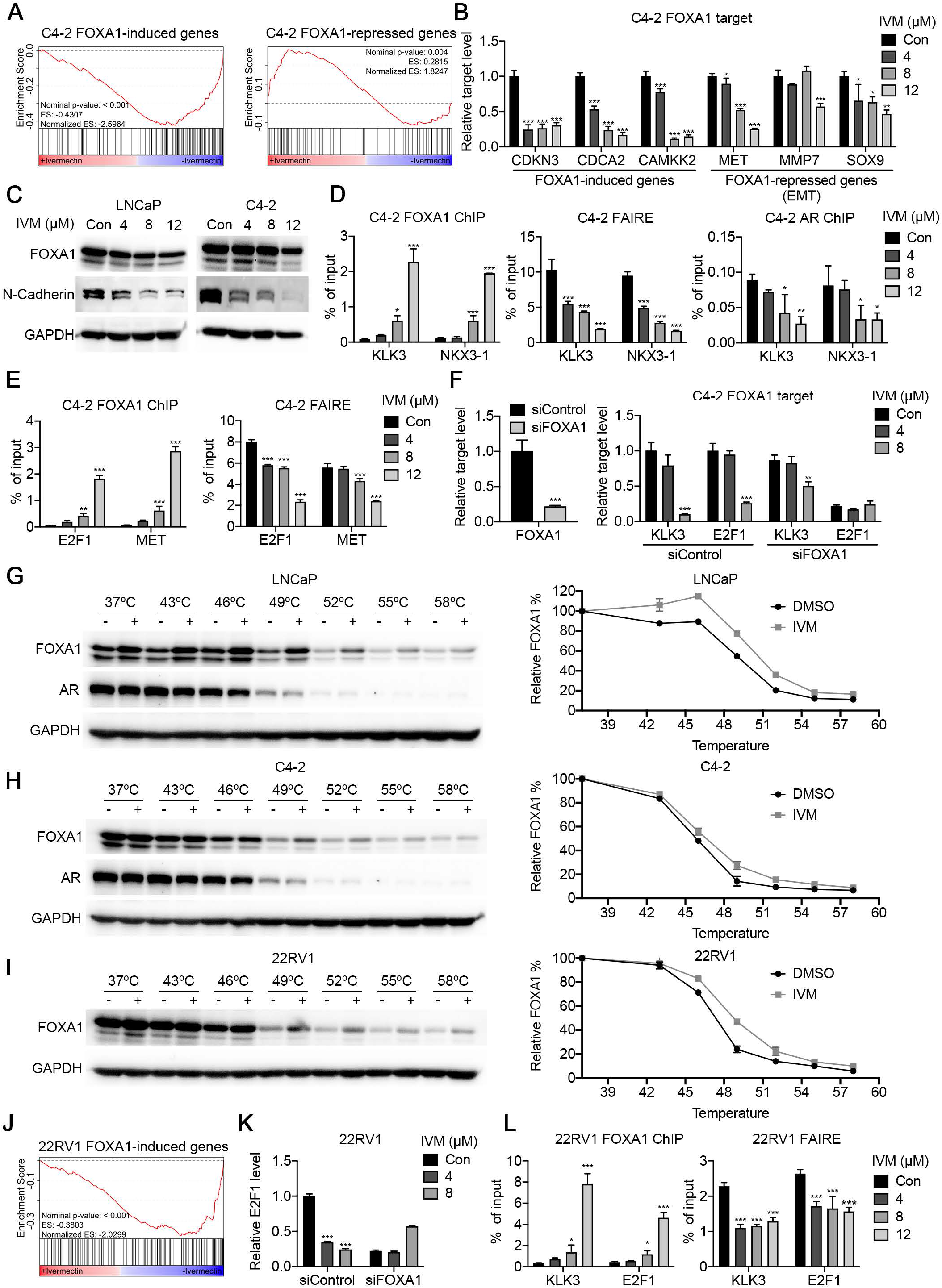
Ivermectin interacted with FOXA1 to block pioneer factor activity. **(A)** GSEA showed that genes induced by FOXA1 were inhibited by ivermectin in C4-2 cells. **(B)** RT-qPCR analysis of FOXA1 induced genes (*CDKN3, CDCA2*, and *CAMKK2*) and FOXA1 repressed EMT associated-genes (*MET, MMP7*, and *SOX9*) in C4-2 cells treated with ivermectin for 48 h. **(C)** Western blot analysis of FOXA1 and N-cadherin in LNCaP and C4-2 cells treated with ivermectin for 48 h. **(D)** ChIP–qPCR analysis for FOXA1 or AR occupancy, and FAIRE–qPCR analysis of chromatin accessibility at a target regulated by AR and FOXA1 (KLK3 and NKX3-1) in C4-2 cells treated with ivermectin. **(E)** ChIP–qPCR analysis for FOXA1 and FAIRE-PCR analysis of chromatin accessibility at a target regulated by FOXA1 (E2F1 and MET) in C4-2 cells treated with ivermectin. **(F)** FOXA1 knockdown impaired the ivermectin-repressed expression of KLK3 and E2F1 genes. mRNA levels were measured 48 h after the implementation of the ivermectin treatment and siRNA transfection by RT-qPCR in C4-2 cells. **(G-H)** Western blots showing thermostable FOXA1 and AR following indicated heat shocks in the presence (+) or absence (−) of 50 μM ivermectin in LNCaP **(G)** and C4-2 **(H)** cells. **(I)** Western blots showing thermostable FOXA1 following indicated heat shocks in the presence (+) or absence (−) of 50 μM ivermectin in 22RV1 cells. **(J)** GSEA showed the inactivation of FOXA1 induced genes in 22RV1 cells after the ivermectin treatment. **(K)** RT-qPCR analysis of FL-AR and ARv7 in 22RV1 cells treated with ivermectin for 48 h. **(L)** ChIP–qPCR analysis for FOXA1 and FAIRE–qPCR analysis of chromatin accessibility at KLK3 and E2F1 in 22RV1 cells treated with ivermectin.

Next, we explored how ivermectin inhibited FOXA1 expression in prostate cancer. ChIP-qPCR and FAIRE-qPCR were performed to explore DNA binding and chromatin accessibility alterations (Jin *et al*, 2014a; Simon *et al*, 2012a). As shown in **Fig. 5D**, the ivermectin treatment increased FOXA1 binding and decreased chromatin accessibility and AR binding on the ARE+FKHD sites of KLK3 and NKX3-1. Similar changes on the ARE+FKHD sites have also been reported by Jin et al(Jin *et al*., 2014a). The authors concluded that excessive FOXA1 enlarges open chromatin regions, which serve as reservoirs that retain AR via abundant half-AREs, thereby reducing AR availability for specific sites. However, we found that although the FOXA1 binding of FKHD-only sites (E2F1 and MET) increased, chromatin was less accessible (**Fig. 5E**). These results were confirmed using the specific ARE+FKHD sites and FKHD-only sites derived from AR and FOXA1 ChIP-seq analysis(Jin *et al*., 2014a) (**Supplementary Fig. S4A**). Increased FOXA1 binding and decreased chromatin accessibility were observed after the ivermectin treatment (**Supplementary Fig. S4B and S4C**). In addition, FOXA1 siRNA transfection alleviated the effect of ivermectin on KLK3 and E2F1 mRNA expression (**Fig. 5F**). Based on these findings, we considered that FOXA1 might be locked on chromatic but unable to loosen the compact chromatin in the presence of ivermectin, thereby inhibiting the transcription of FOXA1 targets, including E2F1 and AR signaling.

Third, the direct binding between FOXA1 and ivermectin was evaluated using CETSA. Our results showed that ivermectin caused the thermal stabilization of FOXA1 in LNCaP and C4-2 cells, but did not affect the thermal stability of AR (**Fig. 5G and 5H**). Increased thermal stability of FOXA1 (**Fig. 5I**) and downregulation of FOXA1 target genes (**Fig. 5J**) were also identified in 22RV1 cells. In line with the results obtained for C4-2 cells, the effect of ivermectin on E2F1 expression was blocked by FOXA1 knockdown (**Fig. 5K**). Meanwhile, increased FOXA1 binding, decreased accessibility, and AR binding were observed in 22RV1 cells (**Fig. 5L**). Thus, ivermectin could target FOXA1 and reduce accessibility in ARV-positive situations.

### The TPP-TR assay revealed that Ku70/Ku80 were additional targets of ivermectin

It is difficult to explain such remarkable cell inhibition after the ivermectin treatment via only targeting FOXA1. Many studies have revealed that ivermectin affects multiple signaling pathways in tumor cells and has been labeled as a “multitargeted” drug(Juarez *et al*., 2018a). Herein, we performed CETSA in a temperature-range thermal proteome profiling (TPP-TR) format, in which protein stability is probed by a mass spectrum, to explore the direct target of ivermectin drugs comprehensively(Berglund *et al*, 2016; Dai *et al*, 2019; Franken *et al*, 2015; Kitagawa *et al*, 2017; Saei *et al*, 2020). The 22RV1 cells were either treated or not with ivermectin (50 μM), and 4,433 complete melting curves were obtained (**Fig. 6A**). The proteins with melting temperature differences (ΔTm) greater than ±3 °C were then screened and subjected to KOBAS KEGG/GO analysis(Jin *et al*, 2014b). We found that targets related to the NHEJ repair pathway (KEGG) and cellular response to gamma radiation (GO) were significantly enriched (**Fig. 6B**). Ku70/Ku80 are important proteins for NHEJ repair. They form heterodimers and recruit DNA-protein kinase catalytic subunit (DNA-PKcs) to the damaged sites that initiate the rejoining of DSB ends(Dietlein *et al*, 2014). The elevated thermal stabilization of Ku70/Ku80 was detected by TPP-TR (**Fig. 6C**) and confirmed by classic CETSA (**Supplementary Fig. S5A**), indicating a direct interaction between ivermectin and the two NHEJ repair proteins. Moreover, we performed CETSA in LNCaP and C4-2 cells. Consistently, the ivermectin treatment increased the thermal stabilization of Ku70/Ku80 (**Fig. 6D and 6E**). Together, these findings show that Ku70/Ku80 are additional direct targets of ivermectin.

**Figure 6.**
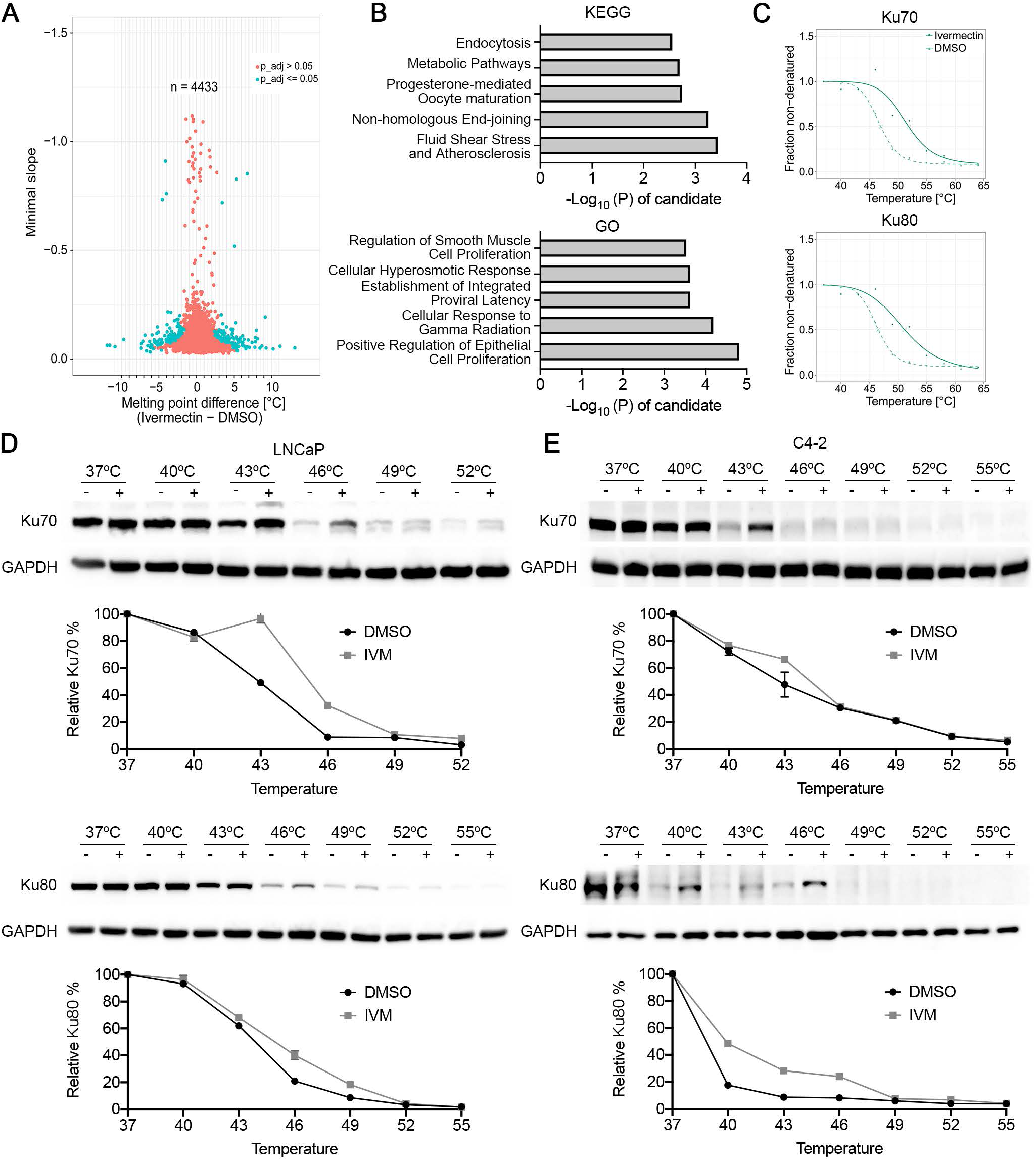
Ivermectin bound to Ku70/Ku80. **(A)** Scatter plot of melting point difference calculated from the ivermectin versus DMSO controls in living 22RV1 cells. Blue circles represent significant melting temperature differences and red circles show all remaining proteins. **(B)** KEGG and GO pathways by KOBAS showed the enrichment pathway of the proteins with the melting temperature difference (ΔTm) more than ± 3 °C. **(C)** Melting curves for Ku70/Ku80 generated from mass spectrum in 22RV1 cells. **(D-E)** Western blots showing thermostable Ku70/Ku80 following indicated heat shocks in the presence (+) or absence (−) of 50 μM ivermectin in LNCaP **(D)** and C4-2 **(E)** cells.

Next, we examined whether the interaction between ivermectin and Ku70/Ku80 influences DNA DSB repair efficiency. The GSEA of gene ontology (GO) gene set revealed ivermectin could decrease the expression of genes associated with DNA repair, with pathway enrichment for DNA recombination repair, DNA recombination, and double strand break repair (**Fig. 7A**). Through western blot, we found ivermectin decreased the expression of homologous recombination (HR) repair pathway executer BRCA1 and Rad51, and inhibited the recruitment of Ku70/Ku80 to the DNA damage site in C4-2 (**Fig. 7B**) and 22RV1(**Fig. 7C**) cells. The BRCA1 and Rad1 were reported as downstream targets of AR(Li *et al*, 2017; Thompson *et al*, 2017) and their mRNA level was consistently decreased after ivermectin treatment (**Supplementary Fig. S5B**). In addition, we evaluated DSB repair efficiency using fluorescent reporter constructs, in which a functional GFP gene was reconstituted following an HR or NHEJ event(Seluanov *et al*, 2010). As expected, the NHEJ and HR repair efficiencies were significantly reduced in ivermectin-treated cells (**Fig. 7D and 7E**).

**Figure 7.**
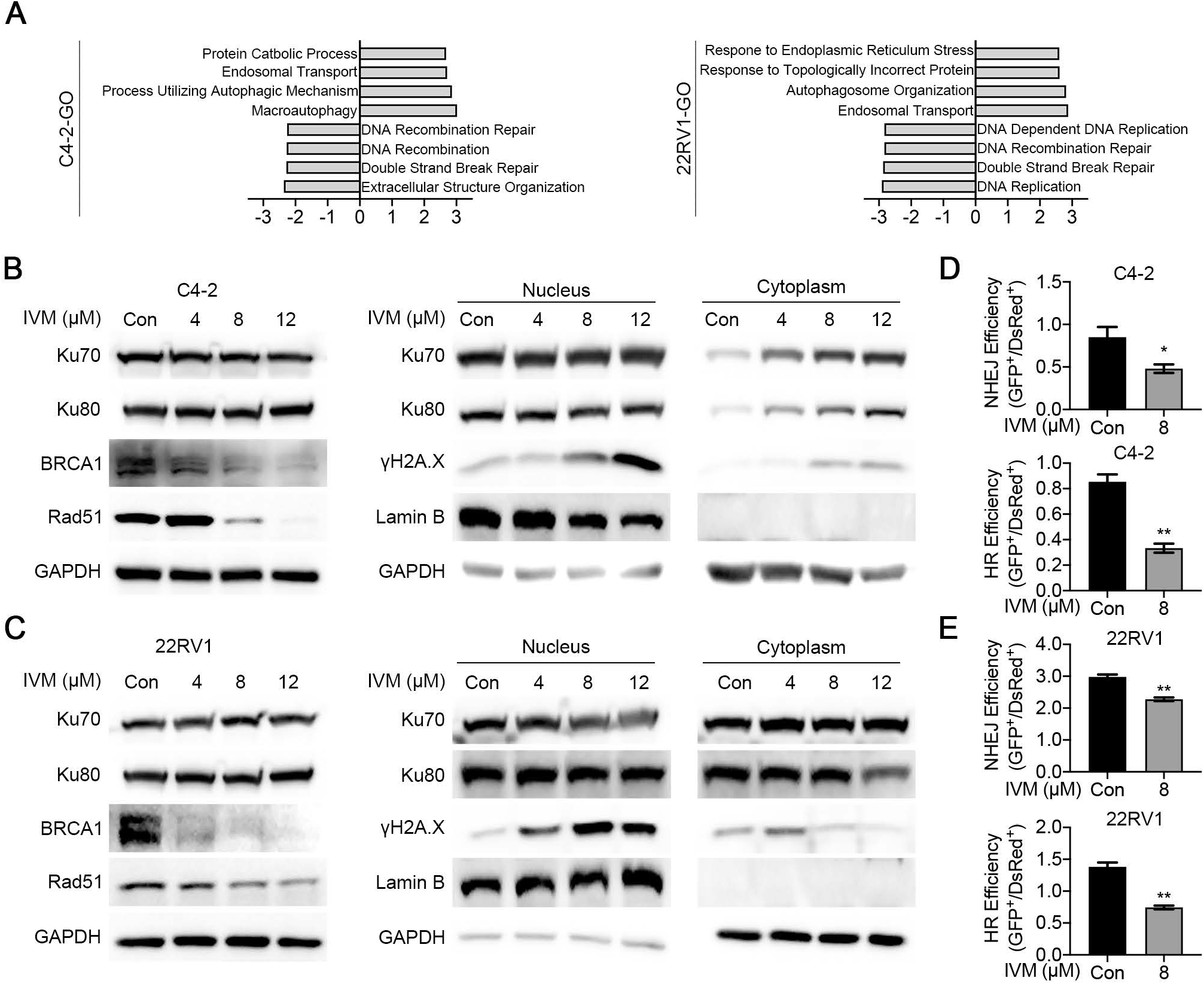
Ivermectin inhibited DSBs repair activity. **(A)** GSEA showed that genes associated DNA damage repair were inhibited by ivermectin in C4-2 and 22RV1 cells. **(B-C)** Western blot analysis Ku70, Ku80, BRCA1, and Rad51 in whole cell lysate or Ku70, Ku80, and γH2A.X in nuclear and cytoplasmic fractions of C4-2 **(B)** and 22RV1 **(C)** cells. Lamin B and GAPDH were probed as nuclear and cytoplasmic loading controls, respectively. **(D-E)** The HR and NHEJ repair efficiencies after the ivermectin treatment were analyzed by flow cytometry using reporter constructs digested in vitro with I-SceI endonuclease, and transfected into C4-2 **(D)** and 22RV1 **(E)** cells as linear DNA. DS-Red was used for transfection control. Repair rate was normalized to DS-Red.

Synthetic lethality has been identified between HR and NHEJ repair(Burdak-Rothkamm *et al*, 2020; Dietlein *et al*., 2014). Based on our results, the inhibition of Ku70/80 recruitment was much more obvious at high doses of ivermectin (12 μM) (**Fig. 7B and 7C**), which is in line with the finding that ivermectin-induced apoptosis was most dramatic at high doses (**Fig. 2B**). We repeated the R1881 experiment with a high-dose ivermectin treatment and found that the R1881 treatment only increased the protein level of Rad51, but exerted no effect on Ku80. The increased HR repair decreased ivermectin-induced cell apoptosis (**Supplementary Fig. S5C**). In AR-negative DU145 cells, CETSA confirmed that ivermectin also bound to Ku70 (**Supplementary Fig. S5D**). The ivermectin treatment did not decrease Rad51 expression, but inhibited the recruitment of Ku70/Ku80 to the DNA damage site (**Supplementary Fig. S5E**). The existence of the HR repair pathway decreased the effect of ivermectin in DU145 cells (**Supplementary Fig. S2B**). Overall, these findings suggest that ivermectin could block NHEJ repair by binding to Ku70/Ku80 and HR repair by downregulating the expression of BRCA1 and Rad51, thereby triggering synthetic lethality in AR-positive prostate cancer cells.

## Discussion

In this study, we reported that ivermectin, an antiparasitic drug, showed promising anticancer activity against prostate cancer progression. Ivermectin was primarily developed for the treatment of onchocerciasis caused by the parasite *Onchocerca volvulus* in poor populations around the tropics(Crump, 2017). Recently, research has shed light on the potential of ivermectin as an antibacterial(Lim *et al*, 2013; Pettengill *et al*, 2012), antiviral(Heidary & Gharebaghi, 2020; Kosyna *et al*, 2015), and anti-cancer agent(Juarez *et al*., 2018a; Tang *et al*., 2020). In particular, owing to its wide margin of clinical safety(De Sole *et al*, 1990), ivermectin is an ideal candidate for drug repurposing and has been listed in the drug repurposing hub established by the Broad Institute(Corsello *et al*, 2017). Our results indicate that ivermectin inhibited dramatically prostate cancer in cell lines representing the hormone-sensitive stage (LNCaP), castration resistance stage (C4-2), and AR variant positive stage (22RV1). In addition, there is controversy regarding the cellular targets of ivermectin, and several alternative action mechanisms have been proposed. To address this issue, we performed an integrated analysis including RNA-seq and TPP-TR to identify the direct targets of ivermectin in prostate cancer. Our data showed that ivermectin could bind to FOXA1 and Ku70/Ku80 directly and inhibit AR signaling, E2F1 expression, and DNA damage repair activity, thereby leading to G0/G1 cell cycle arrest, DNA damage, and trigger synthetic lethality (**Fig. 8**).

**Figure 8.**
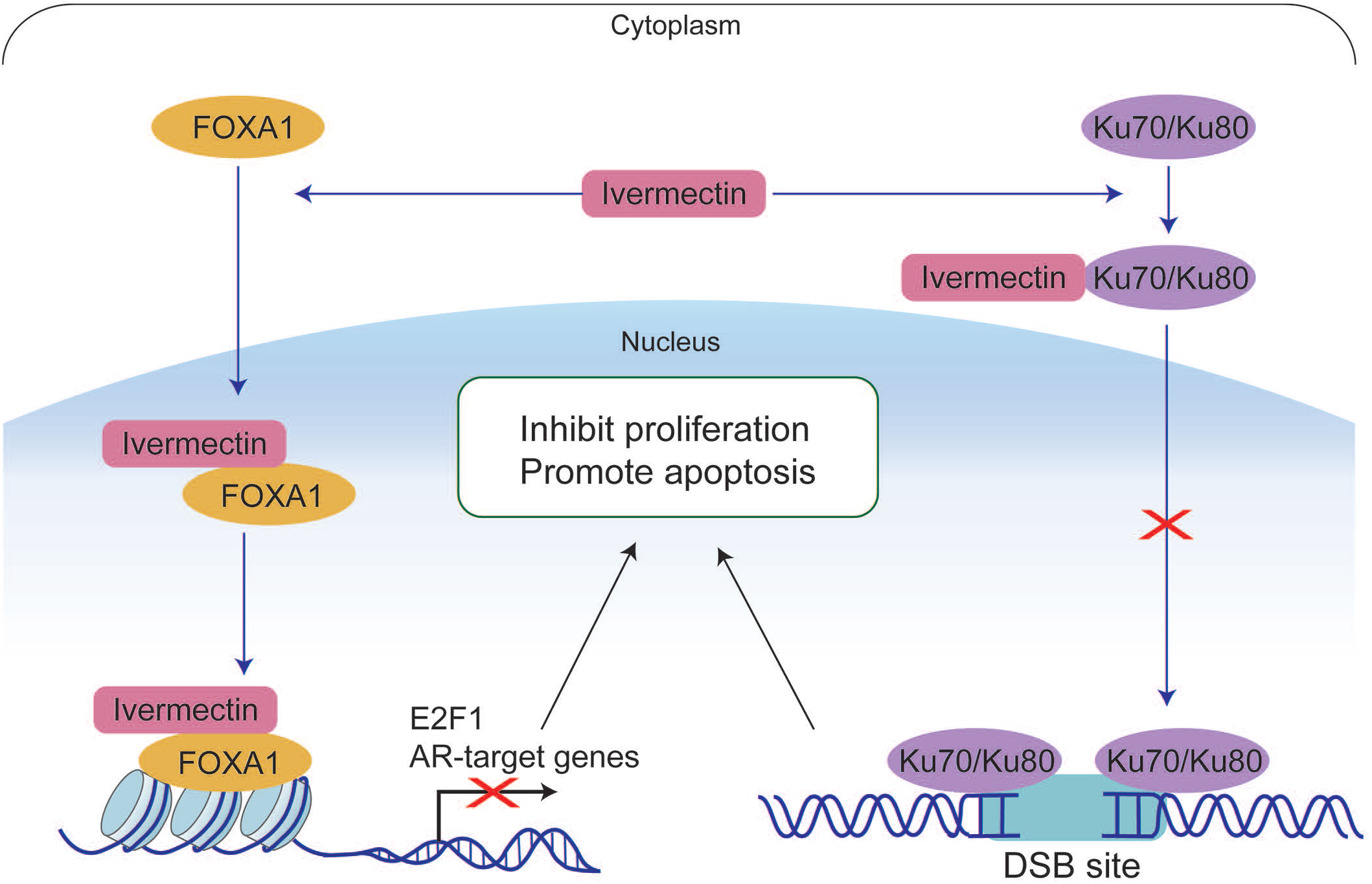
A model for mechanisms of ivermectin inhibiting prostate cancer progression. In PCa, ivermectin could target FOXA1 and Ku70/Ku80 directly and simultaneously. The binding of ivermectin and FOXA1 reduced the chromatin accessibility of AR signaling and E2F1, leading to cell cycle arrest and inhibiting cell proliferation. The binding of ivermectin and Ku70/Ku80 block the recruitment of Ku70/Ku80 to DSB sites. Cooperating with the downregulation of AR regulated homologous recombination repair genes, BRCA1 and Rad51, ivermectin increased intracellular DNA damage level and triggered synthetic lethality.

In our study, ivermectin suppressed AR signaling in CRPC-and ARVs-positive CRPC cells. Targeting the AR signaling axis is the mainstay of prostate cancer therapy. However, stronger inhibition of AR signaling also leads to cancer cell resistance to anti-androgens. In the CRPC stage, AR undergoes changes in expression(Abida *et al*., 2019), structure(Kumar *et al*, 2016) and intracellular localization(Lv *et al*, 2020). These alterations cause AR signaling to re-activate and promote cancer cell proliferation even in the presence of secondary anti-androgens, such as enzalutamide or apalutamide(Fujita & Nonomura, 2019). Herein, we reported that ivermectin could continue blocking AR signaling in both CRPC-and ARV-positive CRPC cells. In contrast to other anti-androgens, ivermectin targets AR through two different mechanisms. First, ivermectin inhibited the AR transcription activity. Our results indicated that ivermectin could block the R1881 induced AR activity in LNCaP and C4-2 cells without significantly reducing AR levels in various prostate cancer cell lines. Second, ivermectin decreased the expression of AR. Nappi et al. proved that ivermectin promotes AR degradation by targeting HSP27(Nappi *et al*, 2020). This combination effect of ivermectin makes it possible to overcome the reactivation of AR induced by overexpression and splice variants. Thus, ivermectin is considered a promising novel antiandrogen for the treatment of enzalutamide-resistant CRPC.

Our research revealed that ivermectin is a novel inhibitor of FOXA1, which blocks the AR and E2F1 signaling pathways. Wang et al. reported that the bromodomain and extraterminal domain (BET) inhibitor JQ1 could independently inhibit FOXA1 and promote prostate cancer invasion(Wang *et al*., 2020). In our study, we identified that ivermectin bound to FOXA1 directly via CETSA. Ivermectin disturbed the pioneering function of FOXA1 and decreased chromatin accessibility. In contrast to JQ1, ivermectin also downregulated the expression of EMT genes, such as *MET, MMP7*, and *SOX9*, and did not induce EMT in prostate cancer. This suggests that the interaction of FOXA1 with ivermectin is different from its interaction with JQ1. Unlike ivermectin, JQ1 did not affect the binding of FOXA1 to its target genes, but inhibited FOXA1 binding to repressors(Wang *et al*., 2020). A recent large-scale integrative genomics study showed that the mutation frequency of FOXA1 is up to 41% in Asian populations(Li *et al*, 2020). The mutations of FOXA1 altered its pioneering activity, perturbing normal luminal epithelial differentiation programs, and prompting prostate cancer progression(Adams *et al*, 2019). Thus, targeting FOXA1 transcription is a very important therapeutic strategy for CRPC treatment. Ivermectin should be further developed as a potent FOXA1 inhibitor.

Our analysis concluded that ivermectin can promote prostate cancer cell death by triggering synthetic lethality. TPP is a high-throughput method for accessing ligand binding in living cells based on the thermal stability of proteins(Franken *et al*., 2015; Savitski *et al*, 2014). In our TPP-TR analysis, Ku70/Ku80 stood out as an additional target of ivermectin. The Ku70/Ku80 heterodimer is the DNA-binding component of DNA-dependent protein kinase, and forms a ring that can specifically bind to exposed broken DNA ends, which is an early and upstream event of NHEJ(Ai *et al*, 2017; Dietlein *et al*., 2014). Our research showed that ivermectin inhibits the recruitment of Ku70/Ku80 to the DNA damage site, thus decreasing the NHEJ repair capacity. In addition, as downstream targets of AR, the HR repair genes BRCA1 and Rad51 could be repressed by AR inhibitors(Li *et al*., 2017; Thompson *et al*., 2017) and were downregulated after the ivermectin treatment. As both are important for DSB repair, the concurrent inhibition of HR and NHEJ could lead to synthetic lethality(Burdak-Rothkamm *et al*., 2020; Dietlein *et al*., 2014). These results were further supported by RNA-seq analysis, as the P53 pathway was highly activated after the ivermectin treatment. Thus, the inhibition of Ku70/Ku80 is an important component of the carcinogenic inhibition of ivermectin in prostate cancer.

## Conclusion

In summary, our results indicate that ivermectin suppressed the AR and E2F signaling pathways, and DNA damage repair capacity by directly targeting FOXA1 and Ku70/Ku80 to inhibit cell proliferation and promote cell apoptosis in prostate cancer. These findings provide insight into both the effects and mechanisms of ivermectin as an anticancer agent. This raises the possibility of broadening the clinical evaluation of ivermectin for the treatment of prostate cancer.

## Methods

### Cell Culture

Prostate cancer cell lines LNCaP, VCaP, and 22RV1 were purchased from Procell Life Science & Technology Co. Ltd. (Wuhan, China). DU145 cell lines were purchased from the American Type Culture Collection (Manassas). C4-2 and LNCaP95 were kindly provided by Dr. Leland WK Chung (Cedars-Sinai Medical Center, Los Angeles, CA) and Dr. Jun Luo (Johns Hopkins University, Baltimore, MD), respectively. VCaP cells were cultured in DMEM (Lonza), while other prostate cancer cells were cultured in RPMI 1640 (Corning). Media were supplemented with 10% FBS (Atlanta Biologicals) or charcoal-stripped FBS (for LNCaP95 cell line) and 1% penicillin/streptomycin. The human prostate primary cells were generated from benign prostatic hyperplasia patient by us previously(Chen *et al*., 2020) and cultured in 50/50 Dulbecco’s modified Eagle’s medium (DMEM)/F12 (Corning), supplemented with 1 μg/mL insulin-transferrin-selenium-X (Invitrogen), 0.4% bovine pituitary extract (Gibco), and 3 ng/mL epidermal growth factor (Gibco). Mycoplasma contamination was tested by PCR.

### MTT assay

Prostate cancer cells and nontumorigenic human prostate primary cells derived from benign prostatic hyperplasia (BPH) patients(Chen *et al*., 2020) were seeded in 96-well plates. The cells were treated with ivermectin (Sellleck) at various concentrations with or without enzalutamide (Sellleck). Cells were then grown for a further 24, 48 or 72 hours. Cell viability was evaluated by the 3-(4,5-dimethylthiazol-2-yl)-2,5-diphenyltetrazolium bromide (MTT, Sigma) assay as described previously(Lv *et al*, 2018).

### Cell cycle analysis

Prostate cells were seeded in 6-well plates and treated with ivermectin at indicated concentrations with or without enzalutamide for 48 hours. Cell cycle distribution was analyzed with PI staining (BD Biosciences). The stained cells were acquired by flow cytometry (BD Biosciences) and analyzed by FlowJo software.

### Cell apoptosis analysis

Prostate cells were seeded in 6-well plates and treated with ivermectin at indicated concentrations for 48 hours. Cell apoptosis was analyzed with FITC Annexin V Apoptosis Detection Kit (BD Biosciences). The stained cells were acquired by flow cytometry and analyzed by FlowJo software. The FITC Annexin V positive and PI negative or FITC Annexin V and PI positive were measured as apoptosis cells.

### Western blot

Prostate cancer cells were lysed by RIPA buffer containing proteasome inhibitor cocktail (Sigma) or performed nucleocytoplasmic fractionation according to the manufacturer’s instructions (G-Biosciences). The samples were analyzed by immunoblotting with primary antibodies to: PARP (Cell Signaling Technology Cat# 9532, 1:1000), cleaved-caspase 3 (Cell Signaling Technology, Cat# 9664, 1:1000), γH2A.X (Cell Signaling Technology Cat# 2577, 1:1000), AR (Santa Cruz Biotechnology Cat# sc-7305, 1:1000), PSA (Cell Signaling Technology Cat# 5365, 1:1000), UBE2C (Cell Signaling Technology Cat# 14234, 1:200), E2F1 (Cell Signaling Technology Cat# 3742, 1:1000), FOXA1 (Cell Signaling Technology Cat# 53528, 1:1000), Ku70 (Cell Signaling Technology Cat# 4588, 1:1000), Ku80 (Cell Signaling Technology Cat# 2180, 1:1000), BRCA1 (Cell Signaling Technology Cat# 9009, 1:1000), Rad51 (Cell Signaling Technology Cat# 8875, 1:1000), Lamin B (Cell Signaling Technology Cat# 13435, 1:1000), GAPDH (Santa Cruz Biotechnology Cat# sc-47724, 1:1000).

### Comet assay

Prostate cancer cells were seeded in 12-well plates treated with ivermectin at indicated concentrations or doxorubicin (DU145 cells, positive control) for 48 hours and collected for DNA damage analysis. DNA damage was quantified using a neutral comet assay by comet assay kit, (Trevigen) following the manufacturer’s protocol.

### Senescence-associated (SA)- β -galactosidase cytochemical staining

Prostate cancer cells were plated into 12-well plates treated with ivermectin at indicated concentrations for 48 hours. Then the cells were fixed in 4% paraformaldehyde and analyzed using an SA-β-Gal kit (Cell Signaling Technology).

### Xenograft tumor model

BALB/c-nude mice (6–8-week-old) were purchased from the Nanfang Hospital and maintained under pathogen-free conditions. The animal use protocol was approved by the Institutional Animal Care and Use Committee in Nanfang Hospital. 22RV1 cells (3 × 10^6^) suspended in 150 μl medium were gently mixed with 150 μl of Matrigel (Corning) and then inoculated subcutaneously in the right flank region of each mouse. Castration was performed after tumor volume reached 300 mm^3^ and treatment was initiated 4 days later. Tumor-bearing BALB/c-nude mice were randomly assigned into two groups and treated with Ivermectin (10 mg/kg, 3 times per week) or vehicle (DMSO:EtOH:Kalliphor/PBS 1:1:8/10). Tumor volume measurements were performed per 3 days and calculated by the formula length × width × depth × 0.52.

### Histology and immunohistochemistry

Tumors were immediately fixed in 10% neutral buffered formalin for 24 hours, progressively dehydrated in solutions containing an increasing percentage of ethanol and embedded into paraffin blocks. Consecutive 4-μm sections were obtained from paraffin blocks. Sections were counterstained with haematoxylin and eosin (H&E), or immunoassayed using antibody to Ki67 (Dako, M7240, 1:100), γH2A.X (Cell Signaling Technology Cat# 80312, 1:200) and PSA (Cell Signaling Technology Cat# 2475, 1:1000) through the immunoperoxidase technique.

### Reverse transcriptase quantitative PCR (RT-qPCR)

Prostate cancer cells were seeded in 6-well plates and treated with ivermectin at indicated concentration for 48 hours. RNA from cells was isolated by TRIzol Reagent (Invitrogen). Reverse transcription was performed with 1 μg RNA using PrimeScript RT reagent Kit (Takara). The cDNA was amplified with gene-specific primers (Supplemental Table 1) and SYBR Premix Ex Taq II kit (TaKaRa). Data were analyzed using a 2^−ΔΔCt^ method.

### RNA-seq and GSEA analysis

C4-2 and 22RV1 cells were treated with 8 or 12 μM ivermectin for 48 hours, and total RNA was extracted by TRIzol Reagent for RNA-Seq analysis. The sequencing data were deposited in the NCBI’s Gene Expression Omnibus (GEO) database (GSE169356). Differentially expressed genes were identified by filtering, with a |log2(FoldChange)| > 1 and *p* adj< 0.05. GSEA was performed using the GSEA Java program (https://www.gsea-msigdb.org/gsea/index.jsp). Normalized enrichment score (NES) and *p* values are shown in the figures.

### ChIP-qPCR

ChIP assays were performed using a Pierce Agarose ChIP Kit (Thermo Fisher Scientific) according to the manufacturer’s protocol. FOXA1 (Abcam, #ab170933), AR (Abcam, #ab108341), and corresponding control IgG antibodies were used. The qPCR assays were carried out using the chromatin samples as prepared above. The primer sequences are listed in Supplemental Table 1.

### Formaldehyde-assisted isolation of regulatory elements qPCR (FAIRE-qPCR)

FAIRE was performed as previously described(Simon *et al*, 2012b). Briefly, ivermectin treated C4-2 and 22RV1 cells were cross-linked by formaldehyde and the chromatin fractions were sheared and extracted identically as for ChIP. Input samples were reverse cross-linked overnight at 65 °C. The FAIRE samples and reverse cross-linked input samples were subjected to two sequential phenol/chloroform/isoamyl alcohol (25/24/1, Sigma) and one chloroform/isoamyl alcohol (24/1, Sigma) extractions. DNA was precipitated with ethanol and treated with RNase A (Invitrogen) for 30 min at 37 °C. Proteins were then digested by proteinase K and DNA-DNA cross-links were reversed by incubating overnight at 65 °C. FAIRE DNA was next purified by Zymo-I spin columns (Zymo) and detected by qPCR assay.

### Cellular thermal shift assay (CETSA)

The CETSA assay was performed as previously described(Lv *et al*., 2020). Prostate cancer cells were treated with 50 μM ivermectin for 1 hour. Cells were suspended in PBS with protease inhibitors, heated at the indicated temperature for 3 minutes. Samples were subjected to 3 freeze-thaw cycles freeze-thaw using liquid nitrogen and centrifuged. Supernatants were collected and detected by western blot.

### siRNA transfection

FOXA1 siRNA and negative control siRNA were synthesized by Ribobio company. Lipofectamine 2000 (Thermo Fisher) was used to transfect these siRNAs into cells.

### Temperature-range thermal proteome profiling (TPP-TR)

Target identification was performed by CETSA coupled with quantitative mass spectrometry using the standard protocol(Franken *et al*., 2015). In brief, 22RV1 cells were treated by 50 μM ivermectin for 1 hour and lysed by combination of freeze/thaw. The supernatant was transferred into microtubes for MS-sample preparation. At least 100 μg of the protein of lowest temperature group (measured with a BCA assay) and equal volume of supernatants was subjected to be labeled by isobaric tandem mass tag 10-plex (TMT10) reagents corresponding to each temperature point. The pooled fractions from each experiment were analyzed using liquid chromatography Easy nLC system (Thermo Fisher Scientific) combined with Q Exactive plus spectrometer (Thermo Fisher Scientific). MS/MS raw files were processed using MASCOT engine (Matrix Science; version 2.6) embedded into Proteome Discoverer 2.2 (Thermo Fisher Scientific). The reference protein database used was the Uniprot_HomoSapiens_20367_20200226 database. The analysis of the protein quantification data from the ivermectin- and DMSO-treated samples is performed using the TR functionality of the TPP package by R.

### DNA damage repair assays

Plasmids containing NHEJ, HR reporter cassettes and pDsRed-N1 as the internal controls were kindly provided by Dr Zhiyong Mao from the School of Life Science and Technology of Tongji University (Shanghai, China)(Seluanov *et al*., 2010). Plasmids containing NHEJ or HR reporter cassettes were linearized by I-SceI restriction enzymes (NEB) and purified using GeneJET PCR purification kit (Thermo Fisher Scientific). Cells were transfected with 0.5 μg of NHEJ reporter construct or 2 μg of HR reporter construct, and 0.1 μg of pDsRed-N1 as internal control by Turbofect (Thermo Fisher Scientific). After 6 hours, the culture medium was replaced by fresh medium containing ivermectin (8 μM). Cells were analyzed by flow cytometry 48 hours after transfection.

### Statistical analysis

Statistical analysis was performed using GraphPad Prism (Version 8.2.1, for macOS, GraphPad Software). Data are presented as the mean ± SD. A parametric t-test (two groups) and one-way ANOVA followed by Dunnett’s multiple-comparisons post-test (for more than two groups) were used when the data sets were found to be normally distributed, with F test comparison of variances or Bartlett’s test of equal variances, respectively. For the data in all figures, statistical significance was set at *P < 0.05, **P < 0.01, ***P < 0.001.

## Abbreviations

AR: Androgen receptor
FOXA1: Forkhead box protein A1
NHEJ: Non-homologous end joining
ADT: Androgen deprivation treatment
CRPC: Castration-resistant prostate cancer
ROS: Reactive oxygen species
DSB: DNA double-strand break
BPH: Benign prostatic hyperplasia
CHX: Cycloheximide
NES: Normalized enrichment score
PDB: Protein Data Bank
PSA: Prostate-specific antigen
AR-FL: Full-length AR
ARVs: AR variants
GSEA: Gene set enrichment analysis
DEGs: Differentially expressed genes
CETSA: Cellular thermal shift assay
EMT: Epithelial mesenchymal transformation
FAIRE-qPCR: Formaldehyde-assisted isolation of regulatory elements qPCR
TPP-TR: Temperature-range thermal proteome profiling
DNA-PKcs: DNA-protein kinase catalytic subunit
HR: Homologous recombination

## Availability of data and material

The sequencing data were deposited in the NCBI’s Gene Expression Omnibus (GEO) database (GSE169356).

## Competing interests

No potential conflict of interest was reported by the authors.

## Funding

This work was funded in part by China Postdoctoral Science Foundation 2020M682800; by Natural Science Foundation of Guangdong Province 2021A1515011023 (to Shidong Lv); by National Natural Science Foundation of China 81872092(to Qiang Wei), NIH grant R50 CA211242 (to LEP), and Department of Urology, University of Pittsburgh (to Zhou Wang).

## Acknowledgements

We would like to thank Dr. Leland W.K. Chung for C4-2 cells and Dr. Jun Luo for LNCaP95 cells.

## Author Contribution Statement

SL and ZeW carried out the cell function and molecular mechanism studies, participated in the sequence analysis, and drafted the manuscript. ML participated in the sequence analysis and performed the statistical analysis. YZ was in charge of the TPP-TR data analysis and participated in figure organization. JZ participated in the ChIP assay. LEP helped revise the manuscript. ZW and QW conceived of the study, and participated in its design and coordination, and helped to draft the manuscript.

## Supplementary Materials

### Supplementary Figure Legends

**Supplementary Figure 1.**
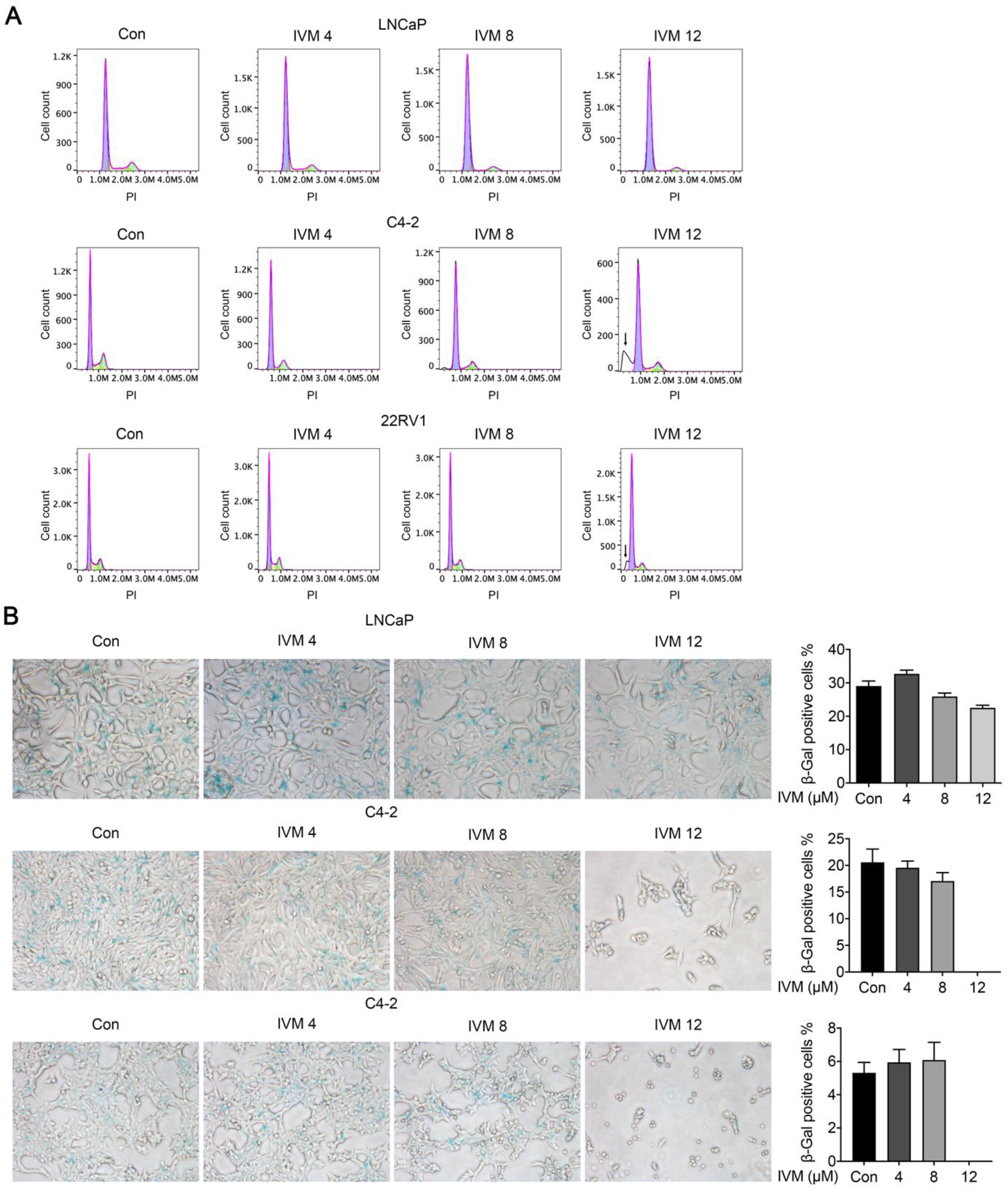
**(A)** Flow cytometry profiling of cell cycle distribution in LNCaP, C4-2 and 22RV1 cells treated with indicated concentrations of ivermectin following PI staining. **(B)** Representative images of SA- β -Galactosidase staining (blue-green) of LNCaP, C4-2 and 22RV1 cells.

**Supplementary Figure 2.**
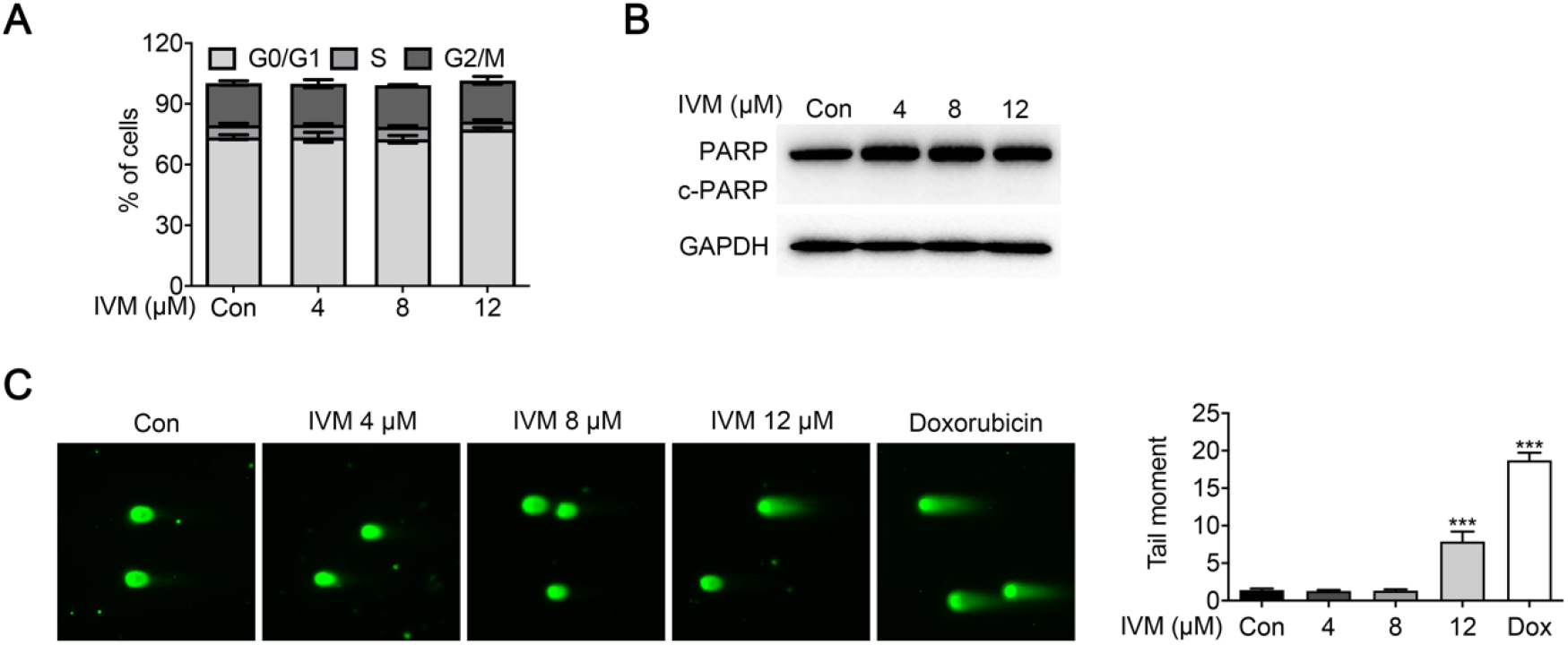
Ivermectin weakly effected AR-negative DU145 cells. **(A)** Ivermectin did not change the cell cycle distribution in DU145 cells treated at 4, 8 and 12 μM for 48 hours. **(B)** Western blot analysis of PARP in cells treated with ivermectin for 48 hours. **(C)** Ivermectin increased DNA damage. DNA fragments were shown as comet images in alkaline gel electrophoresis (Dox: Doxorubicin was used as positive control). The tail moment was used to quantify the DNA damage in the treatment of ivermectin for 48 hours.

**Supplementary Figure 3.**
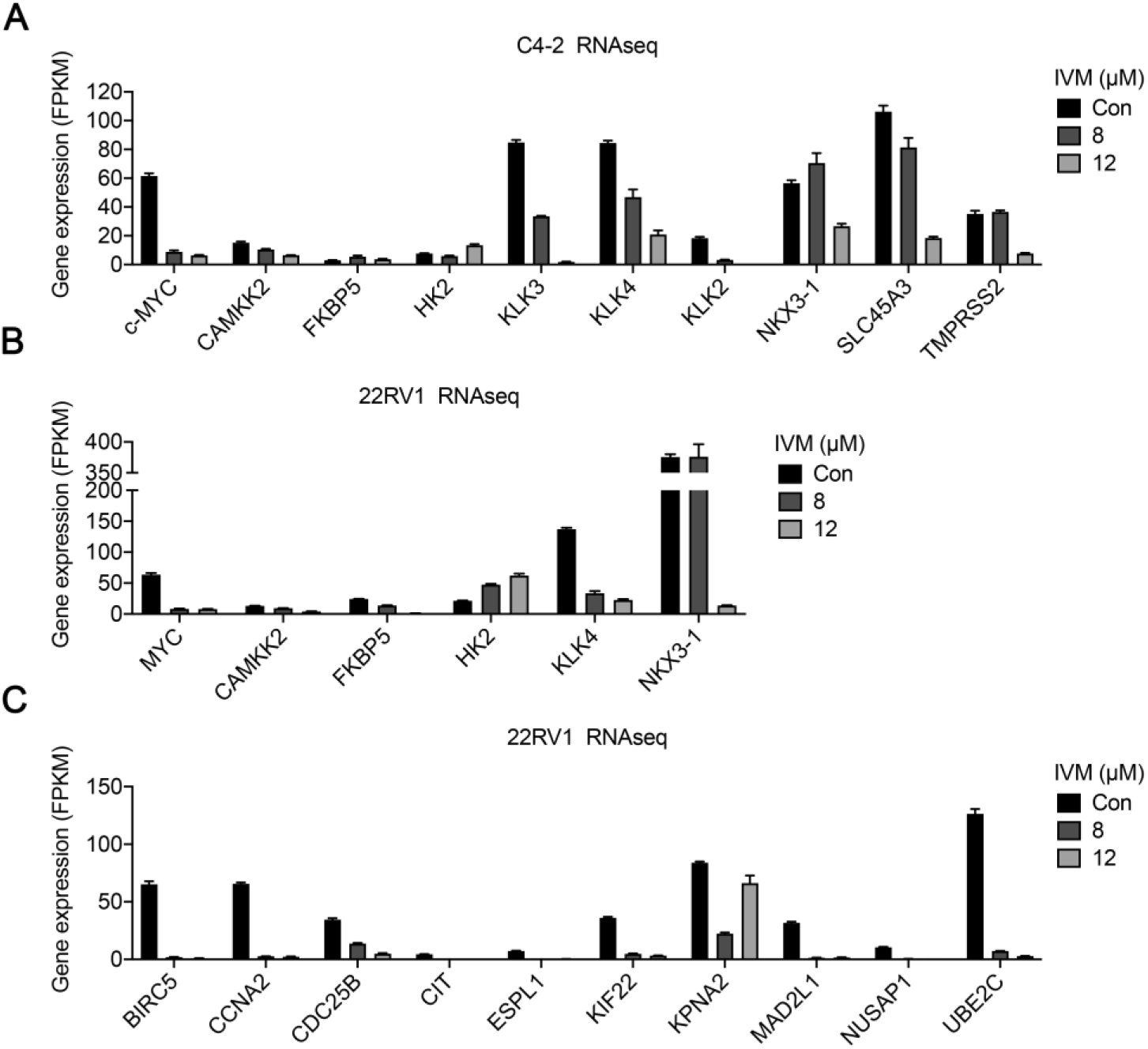
The RT-qPCR verification of differential expression of AR signaling target genes identified by RNA-seq in C4-2 **(A)** and 22RV1 **(B and C)** cells.

**Supplementary Figure 4.**
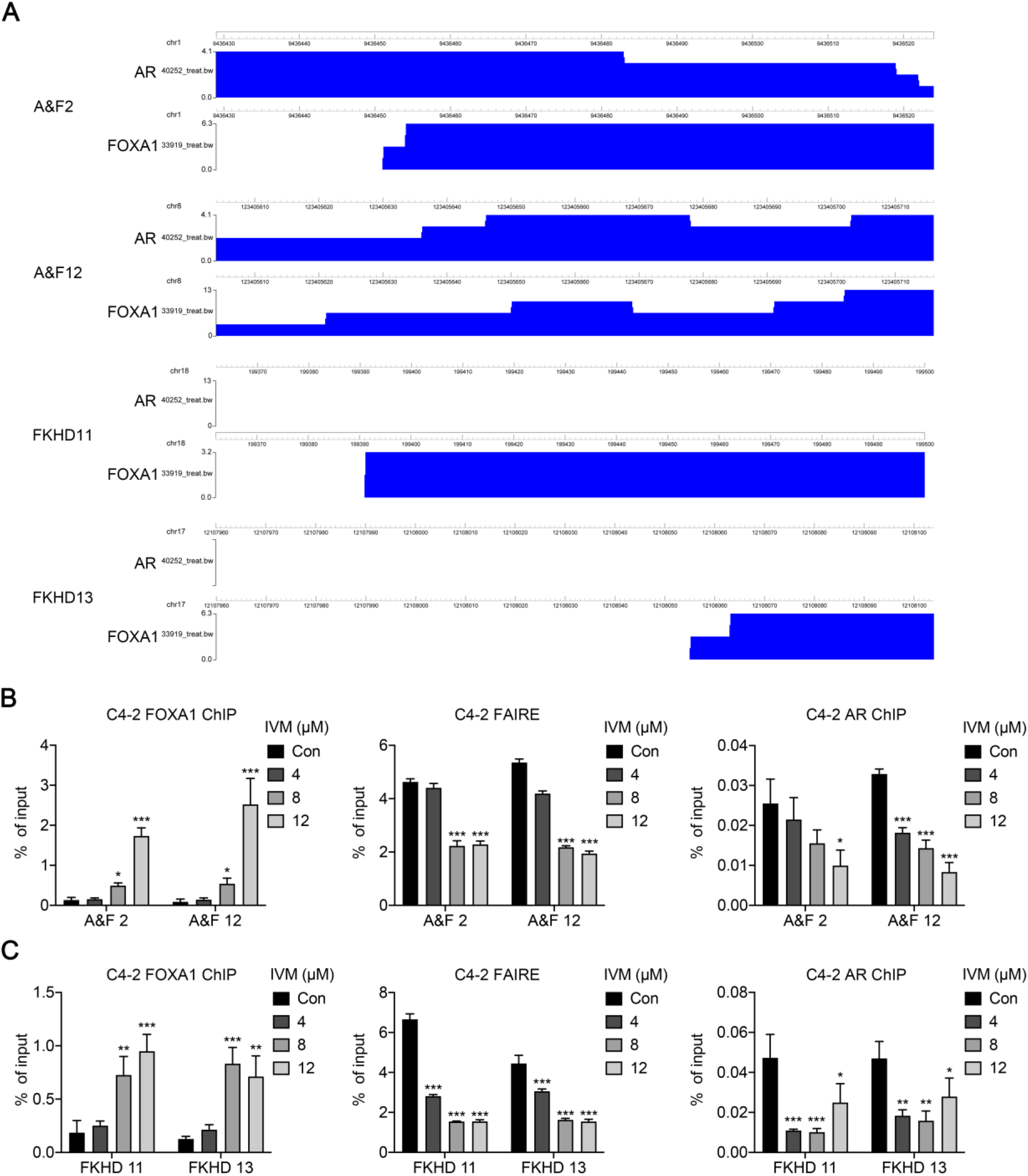
Ivermectin increased the binding of FOXA1 on target sites but decreased the chromatin accessibility. **(A)**. The binding of FOXA1 on ARE+FKHD sites or FKHD only sites by ChIP-seq in LNCaP cells. **(B)** ChIP–qPCR analysis for FOXA1 or AR occupancy, and FAIRE–qPCR analysis of chromatin accessibility at target regulated by AR and FOXA1 in C4-2 cells treated with ivermectin. **(C)** ChIP–qPCR analysis for FOXA1 and FAIRE-PCR analysis of chromatin accessibility at target regulated by FOXA1 in C4-2 cells treated with ivermectin.

**Supplementary Figure 5.**
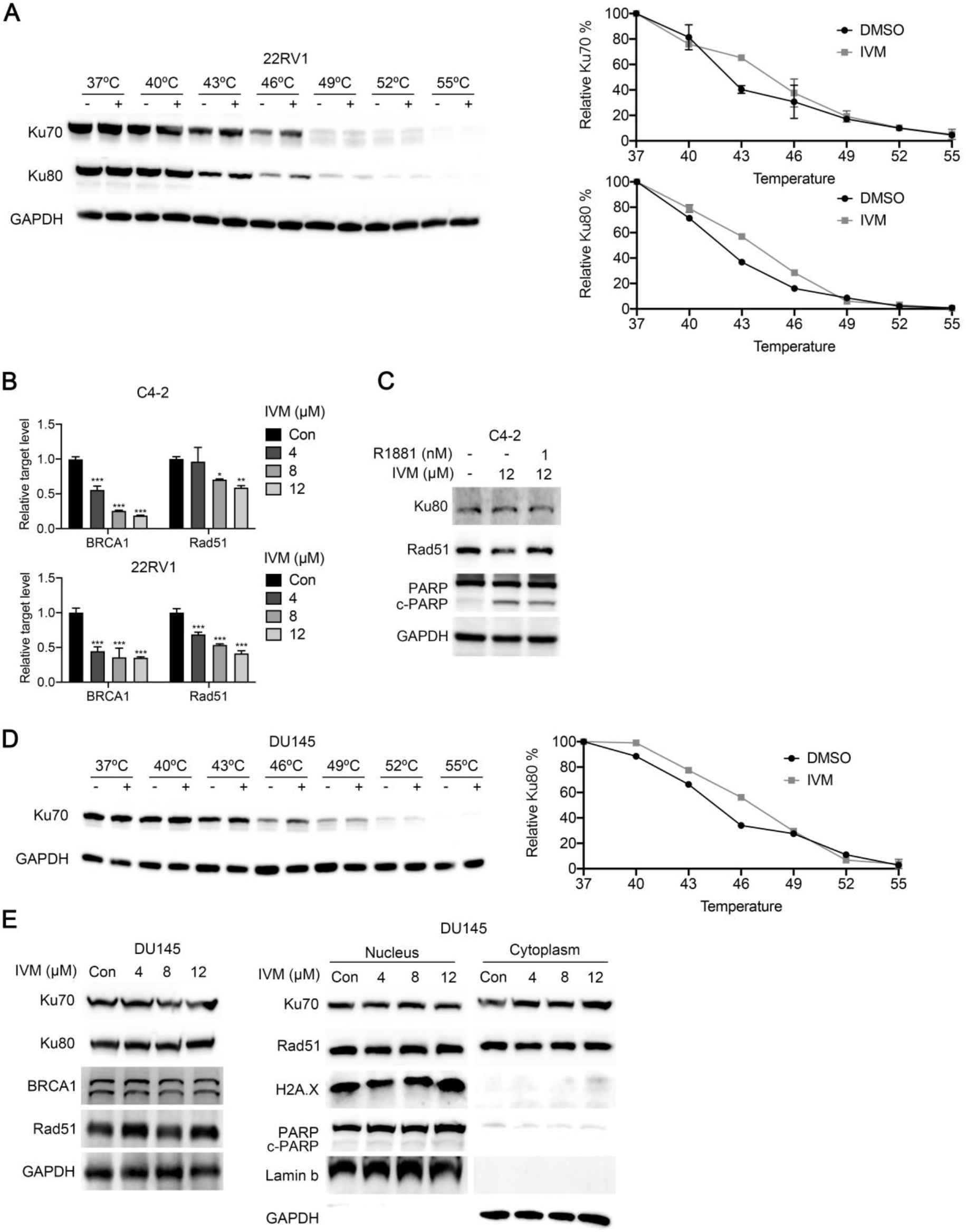
**(A)** Verification of TPP-TR by western blot in 22RV1 cells. **(B)** RT-qPCR analysis of BRCA1 and Rad51 in C4-2 and 22RV1 cells treated with ivermectin for 48 h. **(C)** Western blot analysis of Ku80, Rad51 and PARP and in C4-2 cells after 12 μM ivermectin treatment with or without 1 nM R1881. **(D)** Western blots showing thermostable Ku70 following indicated heat shocks in the presence (+) or absence (−) of 50 μM ivermectin in DU145 cells. **(E)** Western blot analysis of Ku70, Rad51, γH2A.X and PARP in nuclear and cytoplasmic fractions of DU145 cells. Lamin B and GAPDH was probed as nuclear and cytoplasmic loading control, respectively.

**Supplementary Table S1.**
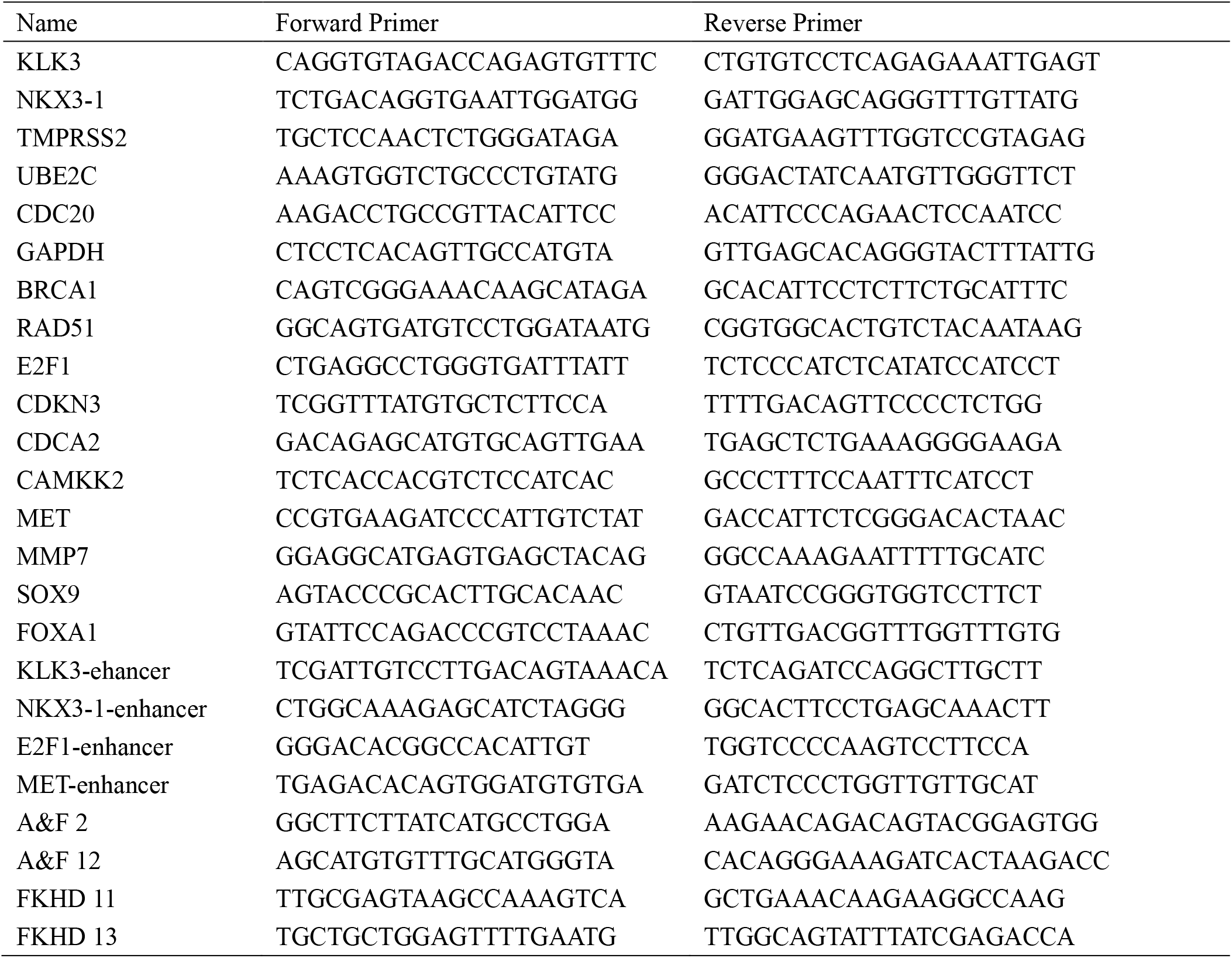
Primer sequence used in RT-qPCR analysis.

## Reference

Abida W, Cyrta J, Heller G, Prandi D, Armenia J, Coleman I, Cieslik M, Benelli M, Robinson D, Van Allen EM et al (2019) Genomic correlates of clinical outcome in advanced prostate cancer. Proc Natl Acad Sci U S A 116: 11428–11436

Adams EJ, Karthaus WR, Hoover E, Liu D, Gruet A, Zhang Z, Cho H, DiLoreto R, Chhangawala S, Liu Y (2019) FOXA1 mutations alter pioneering activity, differentiation and prostate cancer phenotypes. Nature 571: 408–412

Ai J, Pascal LE, Wei L, Zang Y, Zhou Y, Yu X, Gong Y, Nakajima S, Nelson JB, Levine AS (2017) EAF2 regulates DNA repair through Ku70/Ku80 in the prostate. Oncogene 36: 2054–2065

Berglund UW, Sanjiv K, Gad H, Kalderen C, Koolmeister T, Pham T, Gokturk C, Jafari R, Maddalo G, Seashore-Ludlow B (2016) Validation and development of MTH1 inhibitors for treatment of cancer. Annals of oncology 27: 2275–2283

Burdak-Rothkamm S, Mansour WY, Rothkamm K (2020) DNA damage repair deficiency in prostate cancer. Trends in Cancer

Campbell W, Fisher M, Stapley E, Albers-Schonberg G, Jacob T (1983) Ivermectin: a potent new antiparasitic agent. Science 221: 823–828

Carceles-Cordon M, Kelly WK, Gomella L, Knudsen KE, Rodriguez-Bravo V, Domingo-Domenech J (2020) Cellular rewiring in lethal prostate cancer: The architect of drug resistance. Nature reviews Urology 17: 292–307

Chen W, Pascal LE, Wang K, Dhir R, Sims AM, Campbell R, Gasper G, DeFranco DB, Yoshimura N, Wang Z (2020) Differential impact of paired patient-derived BPH and normal adjacent stromal cells on benign prostatic epithelial cell growth in 3D culture. The Prostate 80: 1177–1187

Corsello SM, Bittker JA, Liu Z, Gould J, McCarren P, Hirschman JE, Johnston SE, Vrcic A, Wong B, Khan M (2017) The Drug Repurposing Hub: a next-generation drug library and information resource. Nature medicine 23: 405–408

Crump A (2017) Ivermectin: enigmatic multifaceted ‘wonder’drug continues to surprise and exceed expectations. The Journal of antibiotics 70: 495–505

Dai L, Prabhu N, Yu LY, Bacanu S, Ramos AD, Nordlund P (2019) Horizontal cell biology: monitoring global changes of protein interaction states with the proteome-wide cellular thermal shift assay (CETSA). Annual review of biochemistry 88: 383–408

Davis ID, Martin AJ, Stockler MR, Begbie S, Chi KN, Chowdhury S, Coskinas X, Frydenberg M, Hague WE, Horvath LG (2019) Enzalutamide with standard first-line therapy in metastatic prostate cancer. New England Journal of Medicine 381: 121–131

De Sole G, Dadzie K, Giese J, Remme J (1990) Lack of adverse reactions in ivermectin treatment of onchocerciasis. Lack of adverse reactions in ivermectin treatment of onchocerciasis 335: 1106–1107

Dietlein F, Thelen L, Reinhardt HC (2014) Cancer-specific defects in DNA repair pathways as targets for personalized therapeutic approaches. Trends in genetics 30: 326–339

Dou Q, Chen H-N, Wang K, Yuan K, Lei Y, Li K, Lan J, Chen Y, Huang Z, Xie N (2016) Ivermectin induces cytostatic autophagy by blocking the PAK1/Akt axis in breast cancer. Cancer research 76: 4457–4469

Fang Z, Lin M, Li C, Liu H, Gong C (2020) A comprehensive review of the roles of E2F1 in colon cancer. Am J Cancer Res 10: 757–768

Franken H, Mathieson T, Childs D, Sweetman GM, Werner T, Tögel I, Doce C, Gade S, Bantscheff M, Drewes G (2015) Thermal proteome profiling for unbiased identification of direct and indirect drug targets using multiplexed quantitative mass spectrometry. Nature protocols 10: 1567–1593

Fujita K, Nonomura N (2019) Role of androgen receptor in prostate cancer: a review. The world journal of men’s health 37: 288

Gao S, Chen S, Han D, Barrett D, Han W, Ahmed M, Patalano S, Macoska JA, He HH, Cai C (2019) Forkhead domain mutations in FOXA1 drive prostate cancer progression. Cell research 29: 770–772

Halabi S, Kelly WK, Ma H, Zhou H, Solomon NC, Fizazi K, Tangen CM, Rosenthal M, Petrylak DP, Hussain M (2016) Meta-analysis evaluating the impact of site of metastasis on overall survival in men with castration-resistant prostate cancer. Journal of clinical oncology 34: 1652

Heidary F, Gharebaghi R (2020) Ivermectin: a systematic review from antiviral effects to COVID-19 complementary regimen. The Journal of antibiotics 73: 593–602

Jafari R, Almqvist H, Axelsson H, Ignatushchenko M, Lundbäck T, Nordlund P, Molina DM (2014) The cellular thermal shift assay for evaluating drug target interactions in cells. Nature protocols 9: 2100

Jin H-J, Zhao JC, Ogden I, Bergan RC, Yu J (2013) Androgen receptor-independent function of FoxA1 in prostate cancer metastasis. Cancer research 73: 3725–3736

Jin H-J, Zhao JC, Wu L, Kim J, Yu J (2014a) Cooperativity and equilibrium with FOXA1 define the androgen receptor transcriptional program. Nature communications 5: 1–14

Jin J, Zhang H, Kong L, Gao G, Luo J (2014b) PlantTFDB 3.0: a portal for the functional and evolutionary study of plant transcription factors. Nucleic acids research 42: D1182–D1187

Juarez M, Schcolnik-Cabrera A, Duenas-Gonzalez A (2018a) The multitargeted drug ivermectin: from an antiparasitic agent to a repositioned cancer drug. Am J Cancer Res 8: 317–331

Juarez M, Schcolnik-Cabrera A, Dueñas-Gonzalez A (2018b) The multitargeted drug ivermectin: from an antiparasitic agent to a repositioned cancer drug. American journal of cancer research 8: 317

Kaplun A, Krull M, Lakshman K, Matys V, Lewicki B, Hogan JD (2016) Establishing and validating regulatory regions for variant annotation and expression analysis. BMC genomics 17: 219–227

Kitagawa M, Liao P-J, Lee KH, Wong J, Shang SC, Minami N, Sampetrean O, Saya H, Lingyun D, Prabhu N (2017) Dual blockade of the lipid kinase PIP4Ks and mitotic pathways leads to cancer-selective lethality. Nature communications 8: 1–13

Kodama M, Kodama T, Newberg JY, Katayama H, Kobayashi M, Hanash SM, Yoshihara K, Wei Z, Tien JC, Rangel R (2017) In vivo loss-of-function screens identify KPNB1 as a new druggable oncogene in epithelial ovarian cancer. Proceedings of the National Academy of Sciences 114: E7301–E7310

Kosyna FK, Nagel M, Kluxen L, Kraushaar K, Depping R (2015) The importin α/β-specific inhibitor Ivermectin affects HIF-dependent hypoxia response pathways. Biological chemistry 396: 1357–1367

Kumar A, Coleman I, Morrissey C, Zhang X, True LD, Gulati R, Etzioni R, Bolouri H, Montgomery B, White T (2016) Substantial interindividual and limited intraindividual genomic diversity among tumors from men with metastatic prostate cancer. Nature medicine 22: 369–378

Laing R, Gillan V, Devaney E (2017) Ivermectin–old drug, new tricks? Trends in parasitology 33: 463–472

Li J, Xu C, Lee HJ, Ren S, Zi X, Zhang Z, Wang H, Yu Y, Yang C, Gao X (2020) A genomic and epigenomic atlas of prostate cancer in Asian populations. Nature 580: 93–99

Li L, Karanika S, Yang G, Wang J, Park S, Broom BM, Manyam GC, Wu W, Luo Y, Basourakos S (2017) Androgen receptor inhibitor–induced “BRCAness” and PARP inhibition are synthetically lethal for castration-resistant prostate cancer. Science signaling 10

Lim LE, Vilchèze C, Ng C, Jacobs WR, Ramón-García S, Thompson CJ (2013) Anthelmintic avermectins kill Mycobacterium tuberculosis, including multidrug-resistant clinical strains. Antimicrobial agents and chemotherapy 57: 1040–1046

Lv S, Ji L, Chen B, Liu S, Lei C, Liu X, Qi X, Wang Y, Lai-Han Leung E, Wang H et al (2018) Histone methyltransferase KMT2D sustains prostate carcinogenesis and metastasis via epigenetically activating LIFR and KLF4. Oncogene 37: 1354–1368

Lv S, Song Q, Chen G, Cheng E, Chen W, Cole R, Wu Z, Pascal LE, Wang K, Wipf P (2020) Regulation and targeting of androgen receptor nuclear localization in castration-resistant prostate cancer. The Journal of Clinical Investigation 131: e141335.

Lv S, Wen H, Shan X, Li J, Wu Y, Yu X, Huang W, Wei Q (2019) Loss of KMT2D induces prostate cancer ROS-mediated DNA damage by suppressing the enhancer activity and DNA binding of antioxidant transcription factor FOXO3. Epigenetics 14: 1194–1208

Molina DM, Jafari R, Ignatushchenko M, Seki T, Larsson EA, Dan C, Sreekumar L, Cao Y, Nordlund P (2013) Monitoring drug target engagement in cells and tissues using the cellular thermal shift assay. Science 341: 84–87

Mootha VK, Lindgren CM, Eriksson K-F, Subramanian A, Sihag S, Lehar J, Puigserver P, Carlsson E, Ridderstråle M, Laurila E (2003) PGC-1α-responsive genes involved in oxidative phosphorylation are coordinately downregulated in human diabetes. Nature genetics 34: 267–273

Nappi L, Aguda AH, Al Nakouzi N, Lelj-Garolla B, Beraldi E, Lallous N, Thi M, Moore S, Fazli L, Battsogt D (2020) Ivermectin inhibits HSP27 and potentiates efficacy of oncogene targeting in tumor models. The Journal of clinical investigation 130: 699–714

Pettengill MA, Lam VW, Ollawa I, Marques-da-Silva C, Ojcius DM (2012) Ivermectin inhibits growth of Chlamydia trachomatis in epithelial cells. PLoS One 7: e48456

Rathkopf DE, Smith MR, De Bono JS, Logothetis CJ, Shore ND, De Souza P, Fizazi K, Mulders PF, Mainwaring P, Hainsworth JD (2014) Updated interim efficacy analysis and long-term safety of abiraterone acetate in metastatic castration-resistant prostate cancer patients without prior chemotherapy (COU-AA-302). European urology 66: 815–825

Saei AA, Gullberg H, Sabatier P, Beusch CM, Johansson K, Lundgren B, Arvidsson PI, Arnér ES, Zubarev RA (2020) Comprehensive chemical proteomics for target deconvolution of the redox active drug auranofin. Redox biology 32: 101491

Savitski MM, Reinhard FB, Franken H, Werner T, Savitski MF, Eberhard D, Molina DM, Jafari R, Dovega RB, Klaeger S (2014) Tracking cancer drugs in living cells by thermal profiling of the proteome. Science 346

Seluanov A, Mao Z, Gorbunova V (2010) Analysis of DNA double-strand break (DSB) repair in mammalian cells. JoVE (Journal of Visualized Experiments): e2002

Sharmeen S, Skrtic M, Sukhai MA, Hurren R, Gronda M, Wang X, Fonseca SB, Sun H, Wood TE, Ward R (2010) The antiparasitic agent ivermectin induces chloride-dependent membrane hyperpolarization and cell death in leukemia cells. Blood, The Journal of the American Society of Hematology 116: 3593–3603

Siegel RL, Miller KD, Fuchs HE, Jemal A (2021) Cancer Statistics, 2021. CA: a Cancer Journal for Clinicians 71: 7–33

Simon JM, Giresi PG, Davis IJ, Lieb JD (2012a) Using formaldehyde-assisted isolation of regulatory elements (FAIRE) to isolate active regulatory DNA. Nature protocols 7: 256

Simon JM, Giresi PG, Davis IJ, Lieb JD (2012b) Using formaldehyde-assisted isolation of regulatory elements (FAIRE) to isolate active regulatory DNA. Nat Protoc 7: 256–267

Tang M, Hu X, Wang Y, Yao X, Zhang W, Yu C, Cheng F, Li J, Fang Q (2020) Ivermectin, a potential anticancer drug derived from an antiparasitic drug. Pharmacological Research: 105207

Thompson TC, Li L, Broom BM (2017) Combining enzalutamide with PARP inhibitors: Pharmaceutically induced BRCAness. Oncotarget 8: 93315

Wang L, Xu M, Kao C-Y, Tsai SY, Tsai M-J (2020) Small molecule JQ1 promotes prostate cancer invasion via BET-independent inactivation of FOXA1. The Journal of clinical investigation 130

Zhang C, Wang L, Wu D, Chen H, Chen Z, Thomas-Ahner JM, Zynger DL, Eeckhoute J, Yu J, Luo J (2011) Definition of a FoxA1 Cistrome that is crucial for G1 to S-phase cell-cycle transit in castration-resistant prostate cancer. Cancer research 71: 6738–6748

